# Fucosylated glycoproteins and fucosylated glycolipids play opposing roles in cholera intoxication

**DOI:** 10.1101/2023.08.02.551727

**Authors:** Atossa C. Ghorashi, Andrew Boucher, Stephanie A. Archer-Hartmann, Nathan B. Murray, Rohit Sai Reddy Konada, Xunzhi Zhang, Chao Xing, Parastoo Azadi, Ulf Yrlid, Jennifer J. Kohler

## Abstract

Cholera toxin (CT) is the etiological agent of cholera. Here we report that multiple classes of fucosylated glycoconjugates function in CT binding and intoxication of intestinal epithelial cells. In Colo205 cells, knockout of B3GNT5, the enzyme required for synthesis of lacto- and neolacto-series glycosphingolipids (GSLs), reduces CT binding but sensitizes cells to intoxication. Overexpressing B3GNT5 to generate more fucosylated GSLs confers protection against intoxication, indicating that fucosylated GSLs act as decoy receptors for CT. Knockout (KO) of B3GALT5 causes increased production of fucosylated O-linked and N-linked glycoproteins, and leads to increased CT binding and intoxication. Knockout of B3GNT5 in B3GALT5 KO cells eliminates production of fucosylated GSLs but increases intoxication, identifying fucosylated glycoproteins as functional receptors for CT. These findings provide insight into molecular determinants regulating CT sensitivity of host cells.

## INTRODUCTION

Cholera is a diarrheal disease caused by the bacterium *Vibrio cholerae*.^1^ Cholera toxin (CT) is an AB_5_ protein toxin produced by *V. cholerae* that is the major virulence factor contributing to the disease. AB_5_ family toxins share a common organization comprising a single catalytic A subunit and five copies of the cell surface binding B subunit. The CT holotoxin binds to the surface of epithelial cells in the small intestine via the B subunit (CTB), and is subsequently internalized and retrograde transported through the Golgi to the ER. The catalytic A subunit (CTA) can then dissociate and translocate into the cytoplasm where it ADP-ribosylates Gαs, ultimately leading to ion dysregulation and massive release of fluid into the intestinal lumen.^2^ Though diarrheal symptoms can be easily treated with intravenous fluids or oral rehydration therapy, many fatalities still occur due to difficulties accessing these treatments.

Studies in the 1970s identified a brain ganglioside, GM1, as a high affinity receptor for CT ^3–8^. Gangliosides are a family of glucosylceramide glycosphingolipids (GSLs) characterized by an α2-3 linked sialic acid that is added onto the core GSL structure lactosylceramide (Lac-cer) by the sialyltransferase ST3GAL5^9^. The product of ST3GALT5 action, known as GM3, is subsequently extended with an N-acetylgalactosamine (GalNAc) by the enzyme B4GALNT1 as well as a terminal galactose (Gal) to generate GM1. The interaction between CTB and the GM1 glycan has been structurally characterized^10, 11^, and the affinity depends mainly on the terminal Gal and Neu5Ac (sialic acid) residues, which are recognized in the canonical binding pocket. Although GM1 been demonstrated to be an effective receptor for CT in cell culture, disruption of GM1 biosynthesis in B4GALNT1 KO mice does not result in reduced CT intoxication relative to WT mice.^12^. Furthermore, GM1 expression in human small intestine epithelial tissue is low, making it difficult to accurately detect or quantify^6, 13^. In the course of *V. cholerae* infection, additional GM1 may be produced through the action of *V. cholerae* neuraminidase^6, 14^. However, B4GALNT1 KO mouse intoxication was shown to occur in the absence of *V. cholerae* neuraminidase, indicating that other receptor classes can function in CT intoxication.

In addition to GM1, fucosylated glycoconjugates have been implicated as possible receptors for CT. Epidemiological studies have shown that the severity of cholera symptoms is increased in individuals with the O blood type^15–20^. The ABO blood types are defined by the expression of distinct, fucosylated antigens on the surface of the intestinal epithelium in addition to red blood cell surfaces. CT can directly bind to ABO blood group antigens as well as the related Lewis antigens, which are also fucosylated^12, 21–26^. Structural characterization of CTB revealed a second glycan binding pocket distinct from the canonical GM1-binding pocket.^21, 27^ Fucosylated glycans are recognized in this non-canonical binding pocket. Though the affinity of CTB for fucosylated glycans is orders of magnitude lower than for GM1,^21, 25^ fucosylated glycoconjugates are abundant on the human small intestine epithelial cell surface.^28^ Serendipitously, we found that CTB can bind to fucosylated glycoproteins from intestinal epithelial cell lines and that global reduction of fucosylation reduces CT binding and intoxication^29^. Still, the molecular details of fucosylated structures that regulate host cell intoxication remain incompletely defined.

Here, we report that two distinct classes of fucosylated glycoconjugates regulate CT activity in Colo205 cells. We find that B3GNT5-dependent, fucosylated GSLs act as decoy receptors for CT. Additionally, we observe that disruption of B3GALT5 activity results in increased production of fucosylated glycoproteins, and promotes CT intoxication. Our data are consistent with a model in which the relative expression of fucosylated GSLs and fucosylated glycoproteins control the extent of CT sensitivity in Colo205 cells.

## RESULTS

### CRISPR screen for factors affecting cholera toxin binding to intestinal epithelial cells identifies glycosphingolipid biosynthetic pathway genes

We executed a genome-wide CRISPR/Cas9 KO screen to provide unbiased insight into which glycoconjugates are most important for CTB binding to intestinal epithelial cells (Fig. 1A). Because our goal was to identify cell surface receptors for CTB, it was critical to avoid protease-based methods of cell detachment. Therefore, we selected Colo205 cells, a colorectal cancer cell line that we have used previously in studies of CT, which were most suitable for our screening approach due to their sensitivity to EDTA as a dissociation reagent. We used fluorescence-activated cell sorting (FACS) to collect cells that exhibited altered CTB binding. We collected cells with the lowest 1 % of fluorescence signal (low CTB binding population) as well as the cells with the highest 1 % of fluorescence signal (high CTB binding population).

**Figure 1.**
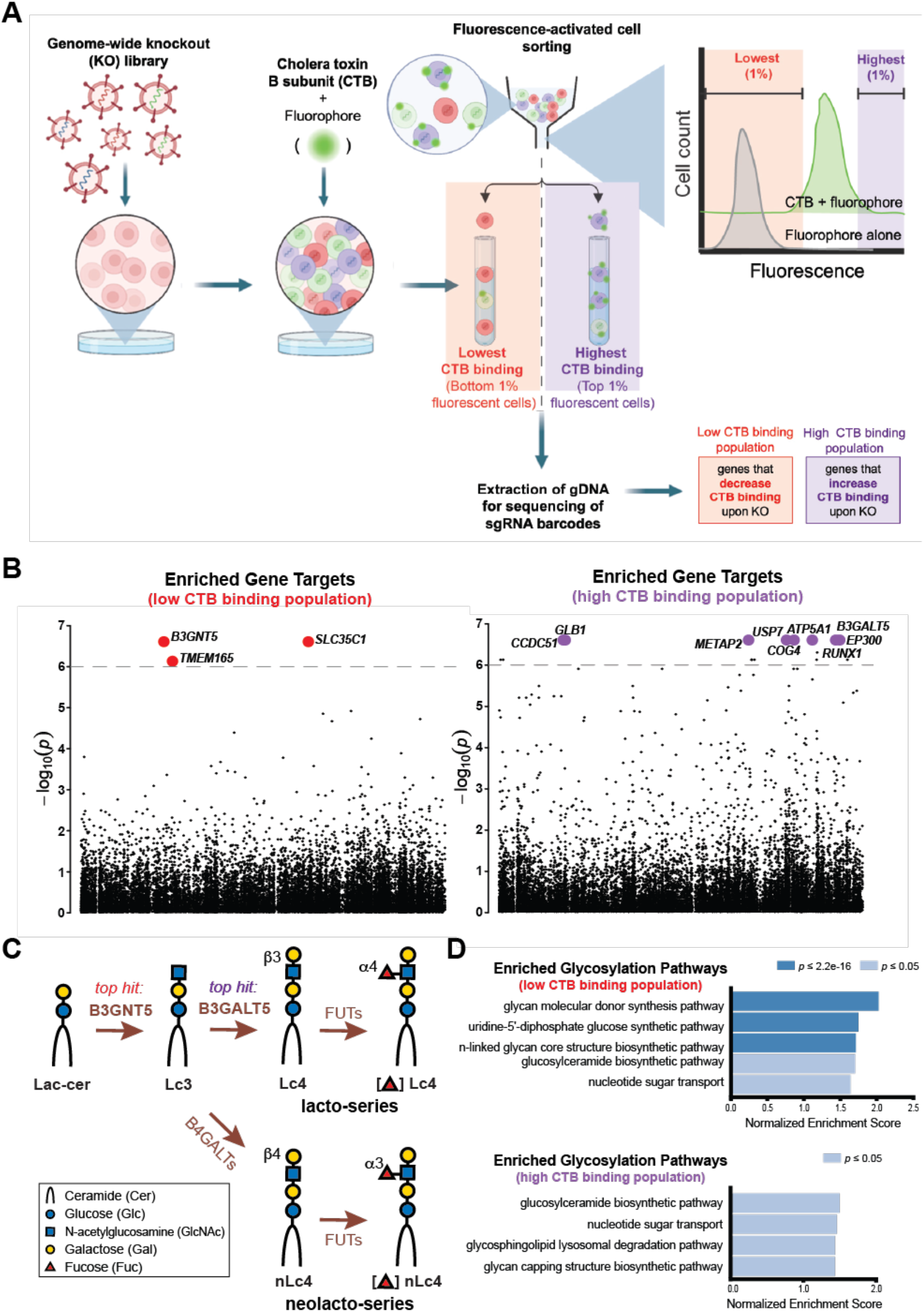
CRISPR screen for factors affecting CTB binding to Colo205 cells identifies glycosphingolipid biosynthetic pathway genes. (**A**) Schematic of genome-wide CRISPR KO screen for mediators of CTB binding to Colo205 intestinal epithelial cells using FACS to collect low versus high CTB binding populations. (**B**) Top sgRNA gene targets (p < 10^−6^) enriched in the low (left panel) and the high (right panel) CTB binding populations relative to the unsorted input population of CRISPR KO library-expressing cells. (**C**) Schematic of lacto- and neolacto-GSL biosynthetic pathway which includes a top enriched sgRNA gene target of both low and high CTB-binding populations. (**D**) GlycoEnzoOnto analysis results identify glycosylation pathways significantly enriched (p < 0.05) in the low (top panel) and high (bottom panel) CTB screen populations.

In the low CTB binding population, three genes (B3GNT5, SLC35C1 and TMEM165) were significantly enriched relative to the input control (p < 10^−6^) (Fig. 1B, left panel). All three have clear links to glycosylation. This is consistent with CTB binding to cell surface glycoconjugates, which is well-documented^3–7, 12, 25, 27, 29–33^. SLC35C1 is a gene that encodes the GDP-fucose transporter^34^. Work from our group and others has established that cell surface fucosylation plays a role in CT binding and intoxication, and CTB has been shown to bind directly to fucosylated glycans^12, 21–23, 25, 27, 29, 31^. Mutations in enriched gene target TMEM165 lead to disrupted Golgi manganese homeostasis and cause a human congenital disorder of glycosylation characterized by pleiotropic glycosylation abnormalities^35^. B3GNT5 encodes an N-acetylglucosaminyltransferase that catalyzes addition of N-acetylglucosamine (GlcNAc) in a β1-3 linkage to galactose (Gal). The preferred substrate onto which B3GNT5 adds GlcNAc is the GSL lactosylceramide (Galβ1-4Glc-ceramide)^36^. The resulting GSL, Lc3 (GlcNAcβ1-3Galβ1-4Glc-ceramide) is the core structure from which all other lacto- and neolacto-series GSLs are generated.

In the high CTB binding population, 17 genes were significantly enriched (p < 10^−6^) (Fig. 1B, right panel). B3GALT5, which encodes a β1-3 galactosyltransferase that catalyzes the addition of Gal onto GlcNAc, was among the gene targets with highest enrichment in this population^37^. B3GALT5 is annotated to function in concert with B3GNT5 in the biosynthesis of lacto-series GSLs (Fig. 1C)^38, 39^ and is highly expressed in Colo205 cells.^40^ Given the abundance of GSL- and other glycosylation-related gene targets in the top hits, we analyzed the CRISPR screening results using GlycoEnzoOnto ontology analysis for pathway identification (Fig. 1D)^41^. Indeed, multiple glycosylation pathways were significantly enriched (p < 0.05) including the nucleotide sugar transport and glucosylceramide biosynthesis pathways, which were enriched in both populations. The related pathways of GSL lysosomal degradation and glycolipid core biosynthesis were enriched in the high and the low CTB binding populations, respectively.

Both the top screen hit from the high CTB binding population (B3GALT5) and the top hit from the low CTB binding population (B3GNT5) have been shown to act on GSLs. Specifically, B3GNT5 catalyzes the formation of Lc3 which can serve as a substrate for B3GALT5 to produce the type 1 structure lactotetraosylceramide (Lc4, Galβ1-3GlcNAcβ1-3Galβ1-4Glc-ceramide) (Fig. 1C). Lc3 can also be used as a substrate by multiple B4GALTs to generate the type 2 structure neolactotetraosylceramide (nLc4, Galβ1-4GlcNAcβ1-3Galβ1-4Glc-ceramide). Since the GSL metabolism pathway was identified as a top hit of the GlycoEnzOnto analysis, we initially hypothesized that CTB binding was mainly dependent upon the relative expression of type 1 versus type 2 GSL structures. To test this, we used CRISPR/Cas9 to knockout B3GALT5 and B3GNT5 individually in Colo205 cells. We obtained two distinct monoclonal populations for each knockout cell line (KO m1 and KO m2).

### Increased binding of CT to B3GALT5 KO cells results in increased intoxication

Since B3GALT5 catalyzes the formation of type 1 LacNAc (Galβ1-3GlcNAc-R) (Fig. 1C), we stained KO cells with antibodies specific for Lewis a (Le^a^), a fucosylated type 1 LacNAc structure (Galβ1-3[Fucα1-4]GlcNAc-R). Le^a^ was completely lost from the surface of B3GALT5 KO cells and restored to control levels by stable overexpression of CRISPR/Cas9-resistant *B3GALT5* (B3GALT5 KO+OE) (Supplementary Fig. 1A). Conversely, B3GALT5 KO+OE cells displayed reduced expression of Lewis x (Le^x^), a fucosylated type 2 LacNAc structure (Galβ1-4[Fucα1-3]GlcNAc-R), as compared to control cells (Supplementary Fig. 1B). These results confirm that B3GALT5 controls expression of the type 1 LacNAc glycans with inverse effects on expression of type 2 LacNAc glycans, consistent with prior _work.38, 39_

Both B3GALT5 KO m1 and B3GALT5 KO m2 exhibited increased CTB binding relative to a scramble sgRNA expressing population, as predicted by the CRISPR screening results (Fig. 2A). The increase in CTB binding to B3GALT5 KO cells was apparent at a range of CTB concentrations (Supplementary Fig. 1C) and overexpression of B3GALT5 reverted CTB binding to below that of the scramble control cells (Fig. 2A). To see if increased CTB binding to B3GALT5 KO cells had an impact on toxin function, we assayed toxin internalization in these cells. We treated both B3GALT5 KO cell lines as well as B3GALT5 KO+OE and control cells with biotin-CTB complexed with streptavidin-saporin (CTB-saporin). Saporin is a cell-impermeable, ribosome-inactivating toxin^42, 43^. Therefore, only active internalization of CTB-saporin can cause saporin-induced toxicity. CTB-saporin treatment caused a larger decrease in cell viability in both B3GALT5 KO m1 and m2 cells as compared to control cells (Supplementary Fig. 1D). Since decreased cell viability is indicative of increased CTB-saporin internalization, this result is consistent with increased CTB binding to functional receptors on B3GALT5 KO cells. Cell viability was restored to control cell levels in B3GALT5 KO+OE cells. Importantly, treatment of all cell lines with unconjugated saporin resulted in minimal cell death, indicating that B3GALT5 KO cell susceptibility to CTB-saporin is not a result of nonspecific toxin internalization (Supplementary Fig. 1E).

**Figure 2.**
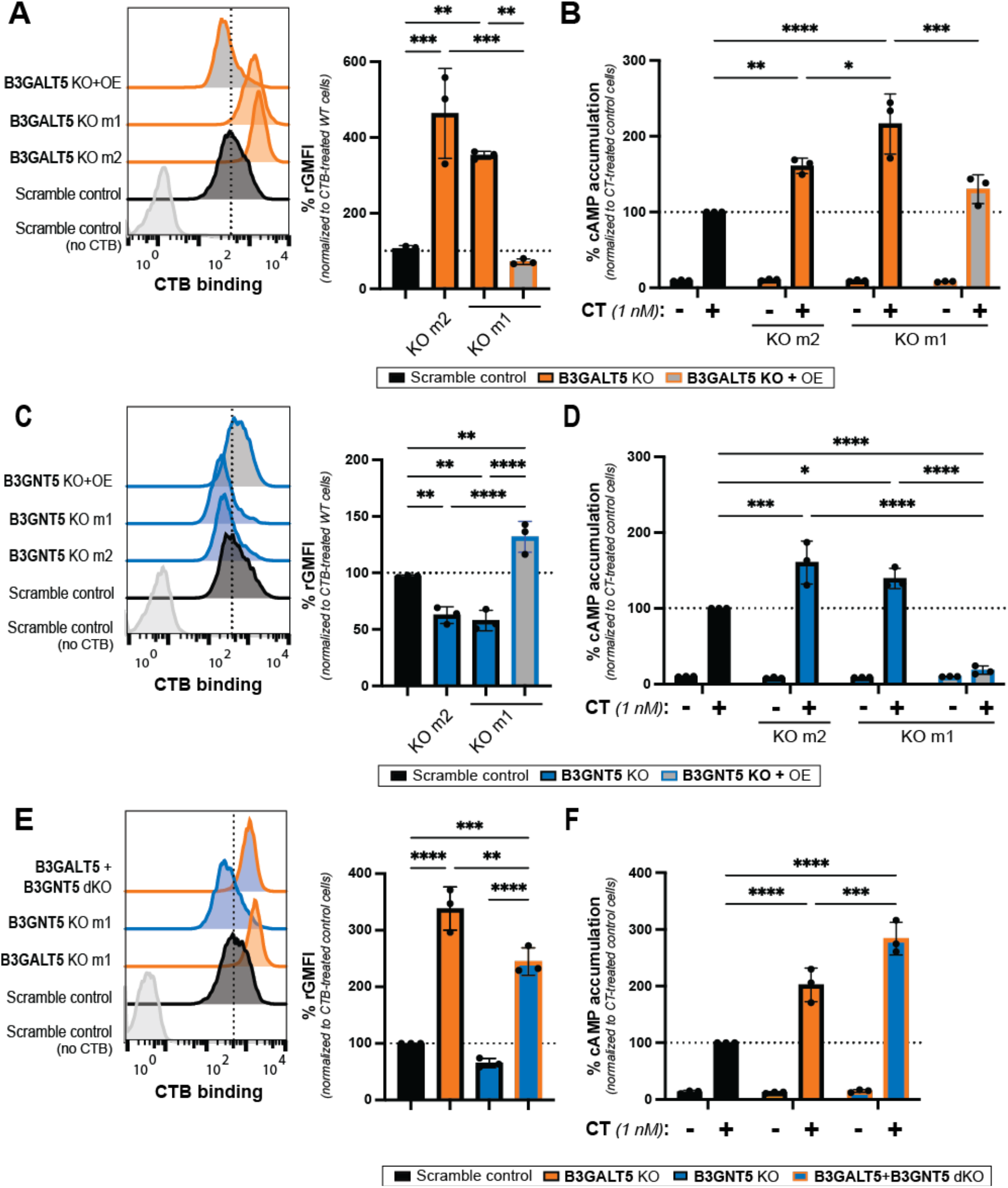
Knockout of B3GALT5 and B3GNT5 results in increased sensitivity to CT. (**A**, **C**, **E**) Representative histograms from the flow cytometry analyses of cell surface binding of CTB (1 μg/mL) to control, KO, KO + OE, and dKO cell lines. Bar graph shows quantification of geometric mean fluorescence index (gMFI) from 3 independent trials, normalized to the maximum APC signal in control cells. Error bars indicate mean ± SD. (**B**, **D**, **F**) Control, KO, KO + OE, and dKO cell lines were incubated for 1.5 h with cholera holotoxin (CTX, 1 nM). Accumulation of cAMP was measured using the cAMP-Glo Max assay. Luminescence values were inversely proportional to cAMP levels. Total amount of plated cells was measured using the Cell Titer-Glo 2.0 assay. Data shown are inverse cAMP values normalized first to the total amount of cells plated for each cell line, then to the maximum signal in control cells. Each datapoint is a biological replicate consisting of 3 averaged technical replicates. Error bars indicate mean ± SD of 3 biological replicates. All statistical analyses were performed by one-way ANOVA with Tukey correction (* indicates p value between 0.01 and 0.05, ** 0.001 and 0.01, *** 0.001 and 0.0001, and **** <0.0001).

To further assess the impact of differential CTB binding and internalization on toxin function, we measured intoxication of B3GALT5 KO m1, m2 and KO+OE cells by CT. Consistent with the binding and internalization results, both B3GALT5 KO m1 and m2 cells were more sensitive to CT, as evidenced by increased cAMP accumulation relative to control cells (Fig. 2B). cAMP accumulation was restored to control cell levels in B3GALT5 KO+OE cells. To evaluate if the observed cAMP accumulation was due to CT intoxication through the canonical pathway, we pre-treated cells with Brefeldin A (BFA), an antiviral agent that disrupts protein transport between ER and Golgi and then added CT. For all cell lines, CT-induced cAMP accumulation was reduced to basal levels (Supplementary Fig. 1F), indicating that a functional secretory pathway is required for CT-induced cAMP accumulation. Incubation of all cell lines with forskolin, a small molecule activator of adenylyl cyclase resulted in similar levels of cAMP accumulation (Supplementary Fig. 1G). This result demonstrated that the genetic knockouts do not perturb cAMP signaling. Therefore, the observed differences in cAMP accumulation are due to differences in cholera toxin receptor expression on the cell surface.

### B3GNT5 KO cells display reduced CTB binding but are sensitized to intoxication

In the B3GNT5 KO cell lines, we observed a decrease in CTB binding, as predicted by the CRISPR screening results (Fig. 2C). The decrease in CTB binding to B3GNT5 KO cells was dose-dependent (Supplementary Fig. 2A). Stably overexpressing CRISPR/Cas9-resistant *B3GNT5* in B3GNT5 KO cells reverted CTB binding to above that of control cells (Fig. 2C). Since B3GNT5 is required for the biosynthesis of both type 1 and 2 GSLs (Fig. 1C), we hypothesized that Le^a^ and Le^x^ expression would be reduced in B3GNT5 KO cells. Indeed, Le^x^ expression was significantly decreased in both B3GNT5 KO cell lines (Supplementary Fig. 2B). However, there was no significant difference in anti-Le^a^ staining between control, B3GNT5 KO or B3GNT5 KO+OE cells lines (Supplementary Fig. 2C). Since B3GNT5 KO cells should not produce lacto-series GSLs, this result suggested that most Le^a^ in Colo205 cells is displayed on glycoconjugates other than lacto-series GSLs.

To see if decreased CTB binding to B3GNT5 KO cells had an impact on toxin function, we assayed toxin internalization in these cells. We treated both B3GNT5 KO cell lines as well as B3GNT5 KO+OE and control cells with CTB-saporin. Although the differences were small, the effects of B3GNT5 KO and OE on CTB internalization were consistent with the effects on CTB binding (Supplementary Fig. 2D and 2E). Next, we measured intoxication of B3GNT5 KO m1, m2 and KO+OE cells by CT. Both B3GNT5 KO cell lines were sensitized to CT relative to control cells (Fig. 2D). This result was unexpected, as binding of CTB to the B3GNT5 KO cells was significantly decreased relative to control cells. Conversely, B3GNT5 KO+OE cells exhibited almost no cAMP accumulation upon incubation with CT, even though CTB binding and internalization were increased in these cells. Importantly, treatment of cells with BFA prior to CT addition caused cAMP accumulation to be reduced to basal levels (Supplementary Fig. 2F). This result indicated that the observed increase in CTX intoxication in the B3GNT5 KO cells was not due to nonspecific internalization. Further, incubation of all cell lines with forskolin resulted in similar levels of cAMP accumulation (Supplementary Fig. 2G). This demonstrated that the significant protection against CT in B3GNT5 KO+OE cells was not due to a general inhibition of cAMP signaling. Thus, the extent of intoxication by CT does not directly correlate with the amount of CTB binding, suggesting that not all CTB binding molecules are functional receptors.

### B3GALT5 KO cells exhibit increased binding to CTB that is independent of B3GNT5 activity

Since B3GNT5 KO cells exhibited increased intoxication despite decreased CTB binding, this implied the existence of decoy receptors on these cells. In contrast, B3GALT5 KO cells exhibited both increased CTB binding and intoxication, which implied the existence of functional receptors on these cells. Thus, we hypothesized that more than one class of CTB-binding glycoconjugates exist in Colo205 cells. To directly test the role of lacto- and neolacto-series GSL biosynthesis in B3GALT5 KO cells, we used CRISPR/Cas9 to knockout B3GNT5 in the B3GALT5 KO m1 cell line (B3GALT5+B3GNT5 dKO). A monoclonal population of B3GALT5+B3GNT5 dKO cells exhibited decreased CTB binding relative to the parental B3GALT5 KO m1 cell line (Fig. 2E). However, binding was still significantly higher than in control and B3GNT5 KO m1 cells. Furthermore, Le^x^ expression in B3GALT5+B3GNT5 dKO cells was not significantly decreased relative to the parental B3GALT5 KO m1 population (Supplementary Fig. 3A). Together, these results confirmed that increased Le^x^ expression as well as increased CTB binding of B3GALT5 KO cells were independent of the lacto- and neolacto-series GSL biosynthetic pathway.

Since CTB binding is not always indicative of toxin function, we also assayed toxin internalization by B3GALT5+B3GNT5 dKO cells. We treated these cells alongside control and parental B3GALT5 KO m1 cells with CTB-saporin, and observed only partial rescue of the increased internalization observed in B3GALT5 KO m1 parental cells (Supplementary Fig. 3B). While this is consistent with the decreased CTB binding observed by flow cytometry, we noticed that B3GALT5 KO m1 cells exhibited increased internalization of unconjugated saporin relative to control and B3GALT5+B3GNT5 dKO cells (Supplementary Fig. 3C). To further characterize the effect of lacto- and neolacto-series GSL biosynthesis on CT function in B3GALT5 KO cells, we measured CT intoxication of B3GALT5+B3GNT5 dKO alongside control and B3GALT5 KO m1 cells. Strikingly, B3GALT5+B3GNT5 dKO cells exhibited increased cAMP signal upon treatment with CT, even relative to the sensitized B3GALT5 KO m1 parental cell line (Fig. 2F). Intoxication could be blocked by pretreatment with BFA (Supplementary Fig. 3D), indicating that the canonical intoxication pathway is required, as observed in single KO cells. Increased intoxication of dKO cells was not due to changes to cAMP signaling, as evidenced by similar cAMP accumulation across cell lines when treated with forskolin (Supplementary Fig. 3E) Together, these data support our proposal that B3GALT5 KO cells produce functional receptors for CT while B3GNT5 KO cells lose production of decoy receptors for CT.

### Glycosphingolipids act as decoy receptors for cholera toxin

Because loss of B3GNT5 activity sensitized cells to CT, we considering the possibility that B3GNT5 KO might shunt precursor GSLs to the biosynthesis of other GSLs such as GM1. We therefore assessed the GSL composition of B3GNT5 KO as well as B3GALT5 KO and control cells by nSI-MS/MS (Fig. 3A and Supplementary Fig. 4A, 4B, and 4C). While GM1 was not detected in any of the cell lines, control cells expressed extended, fucosylated GSLs, which were identified by accurate mass and MS/MS fragmentation analysis (Supplementary Fig. 4D). Fucosylated glycolipids were not detected in either B3GNT5 KO or B3GALT5 KO cells.

**Figure 3.**
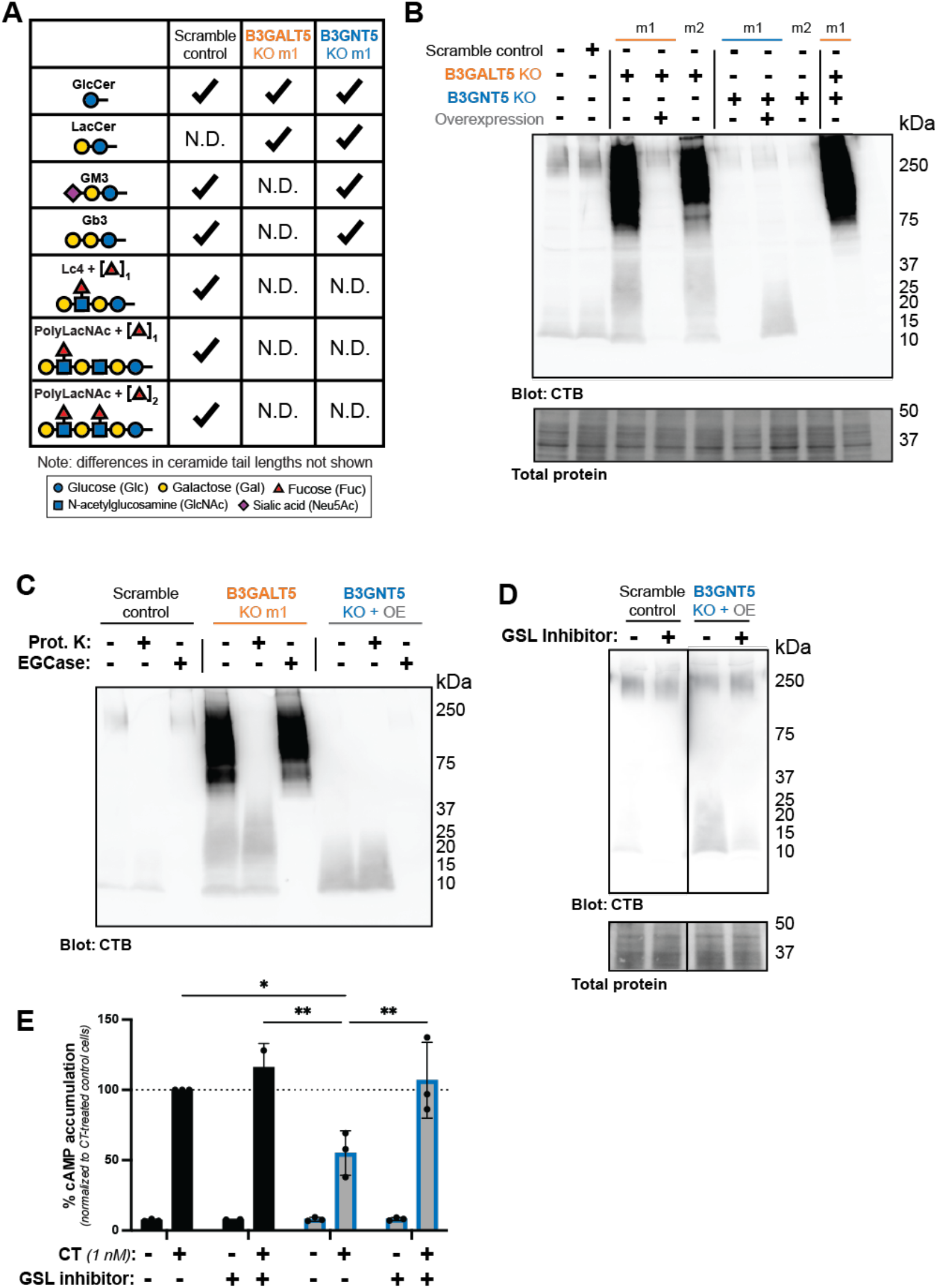
GSLs are decoy receptors for CT. (**A**) Table of glycan headgroups detected by MS analysis of glycolipids extracted from control, B3GALT5 KO m1 and B3GNT5 KO m1 cells. “N.D.” indicates that the glycan headgroup was not detected in the corresponding cell line. (**B**) Lectin blot with CTB-biotin of lysates from control, B3GALT5 KO m1, B3GNT5 m1, respective KO+OE, and B3GALT5+B3GNT5 dKO cells. Data shown are a single representative trial of 3 independent biological replicates. (**C**) Lectin blot with CTB-biotin of control, B3GALT5 KO m1, and B3GNT5 KO+OE cell lysates treated for 16 h with proteinase K, endoglycoceramidase, or a vehicle control. Data shown are a single representative trial of 3 independent biological replicates. (**D**, **E**) Control, B3GALT5 KO m1, and B3GNT5 KO+OE cells were treated with P4 inhibitor of glycosphingolipid biosynthesis for 72 h Treated cells were then lysed and subjected to ligand blot analysis with CTB-biotin (**D**) or incubated for 1.5 hrs. with CT (1 nM) for analysis of cAMP accumulation (**E**). Data shown are inverse luminescence values normalized to total amount of cells then to the maximum signal in control cells. Each datapoint is a biological replicate consisting of 3 averaged technical replicates. Error bars indicate mean ± SD of 3 biological replicates. Statistical analysis was performed by two-way ANOVA with Tukey correction (* indicates p value between 0.01 and 0.05 and ** indicates p value between 0.001 and 0.01).

**Figure 4.**
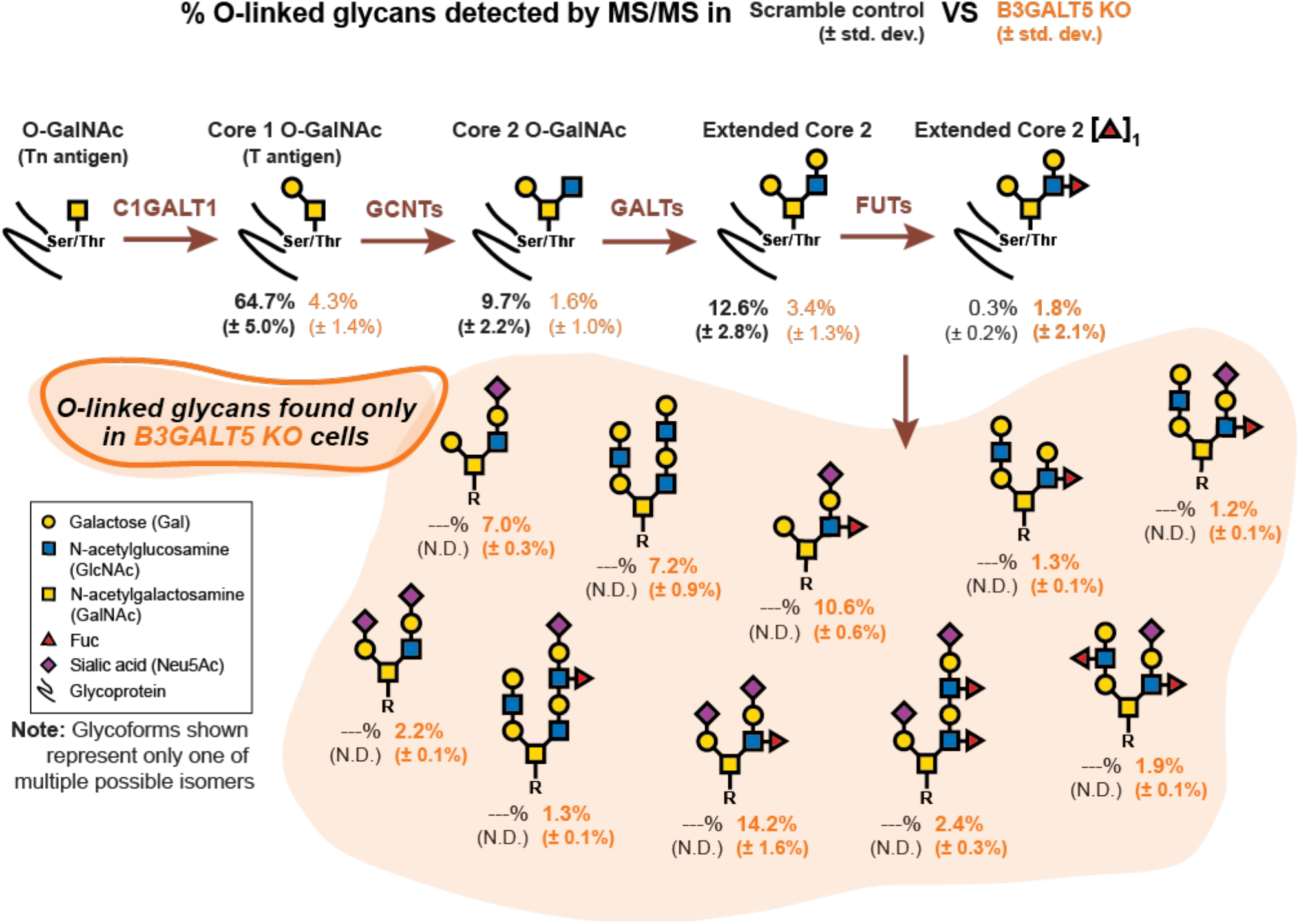
B3GALT5 KO cells exhibit increased fucosylation on O-linked glycoproteins. Schematic of O-linked glycan biosynthesis. Relative enrichment is included below each structure as detected by LC-MS/MS of glycoproteins from control and B3GALT5 KO m1 cells. “N.D.” indicates glycans that were not detected in corresponding cell line. Glycans that were detected in B3GALT5 KO m1 cells but not in control cells are highlighted by orange background.

Since no additional GSLs were detected in B3GALT5 KO cells, we hypothesized that these cells might display increased CTB binding due to altered protein glycosylation. To characterize differential recognition of glycoproteins by CTB, we performed lectin blot analysis of lysates, probing with biotin-CTB. We observed increased CTB recognition of multiple high (> 50 kDa) and low (< 37 kDa) apparent molecular weight (MW) species in B3GALT5 KO m1 and m2 cell lysates as compared to control cell lysates (Fig. 3B). These bands were not observed in B3GALT5 KO+OE rescue cell lysates nor in those of B3GNT5 KO cell lines. Only one band (∼10 kDa apparent MW) was recognized by CTB in lysates of B3GNT5 KO+OE cells. Since these cells were significantly protected against CT intoxication while B3GNT5 KO cells were sensitized to intoxication, we hypothesized that this band might consist of decoy receptors for CT synthesized by B3GNT5. The low apparent MW (∼10 kDa) of the band indicated that the putative decoy receptor might be a GSL. We therefore treated lysates with proteinase K or endoglycoceramidase (EGCase) overnight prior to lectin blot analysis. Proteinase K is a non-specific protease that digests all proteins, while EGCase is an enzyme that cleaves the glycan headgroup from GSLs. We also included B3GALT5 KO cells in this analysis since additional low apparent MW species (∼20 kDa) were detected in these lysates. Indeed, both low apparent MW bands (10 and 20 kDa) recognized by CTB were sensitive to EGCase but not to proteinase K, confirming that these are indeed GSLs (Fig. 3C). To validate our method, we made use of GM1, a GSL that is also a known receptor of CTB^3–7, 30^. Purified GM1 was readily detected by CTB lectin blot and treatment with EGCase completely abrogated detection (Supplementary Fig. 5A). Importantly, GM1 migrated at a lower apparent MW than the species detected in Colo205 cell lysates.

To assess whether GSLs function as decoy receptors for CT, we treated cells with P4, an inhibitor of GSL biosynthesis ^44^. We performed a ligand blot analysis to ensure the specific GSLs of interest were decreased. As expected, cells treated with P4 displayed reduced expression of the low MW GSLs (Fig. 3D). We then measured intoxication of P4-treated cells by measuring cAMP accumulation. When treated with P4, B3GNT5 KO+OE cells were no longer protected against CT, as seen by cAMP accumulation returning to levels similar to those of control cells (Fig. 3E and Supplementary Fig. 5B and 5C). This result is consistent with the hypothesis that the ∼10 kDa GSL species is a decoy receptor.

**Figure 5.**
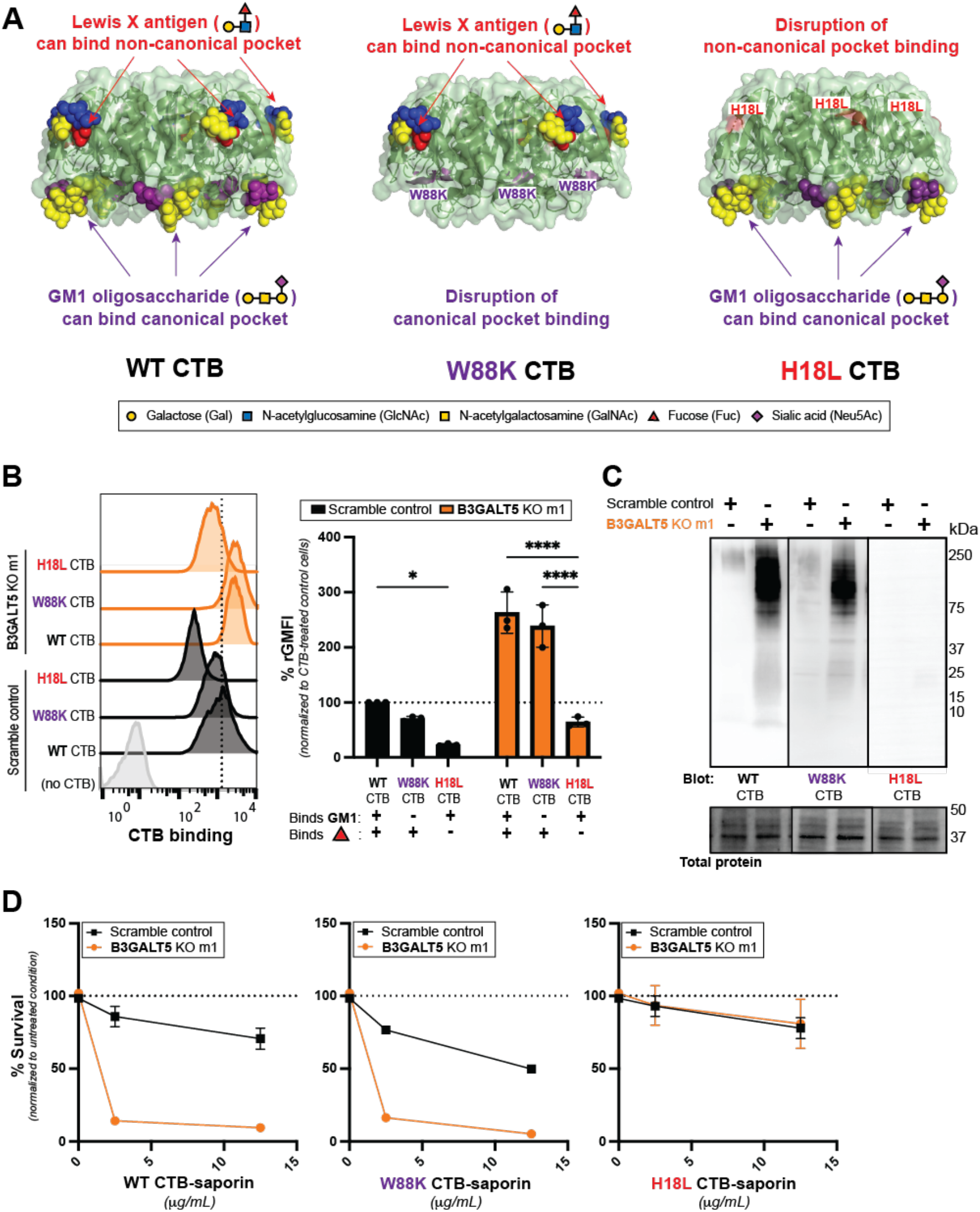
CTB binding to Colo205 cells depends on the non-canonical glycan binding pocket. (A) Structure of CTB pentamer illustrating the canonical and non-canonical binding pockets. Protein and GM1 coordinates from ref. ^11^ and Le^x^ coordinates from ref. ^27^. CTB structures were aligned using Pymol. The locations of the W88K point mutation in the canonical binding pocket and the H18L point mutation in the non-canonical binding pocket are highlighted. (B) Representative flow cytometry histograms of control and B3GALT5 KO m1 cells incubated with WT CTB, W88K CTB or H18L CTB (1 μg/mL). Data shown are a single representative trial from 3 independent experiments. Bar graphs show the quantification of gMFI from the 3 independent trials of normalized to the maximum signal in control cells. Error bars indicate mean ± SD. Statistical analyses were performed by two-way ANOVA with Tukey correction, ∗ indicating *p* value between 0.01 and 0.05, ∗∗ indicating *p* value between 0.001 and 0.01, and ∗∗∗∗ indicating *p* value ≤0.0001. (**C**) Lysates from B3GALT5 KO m1 and control cells were analyzed by ligand blot, probing with either WT or mutant CTB-biotin. Data shown are a single representative trial of 3 independent biological replicates. (**D**) B3GALT5 KO m1 and control cells were incubated for 72 h with increasing concentrations of WT, W88K, or H18L CTB-saporin. Cell survival was measured using the Cell Titer-Glo 2.0 assay. Data shown are luminescence values normalized to the maximum signal for each cell type. Each datapoint is a biological replicate consisting of 3 averaged technical replicates. Error bars indicate mean ± SD of 2 biological replicates.

### CTB recognizes B3GALT5 KO cell receptors through fucose-binding pocket

Since B3GALT5+B3GNT5 dKO cells were more sensitized to CT than other cell lines (Fig. 2F), we hypothesized that they produce functional CT receptors. Lectin blot analysis revealed that B3GALT5 KO results in increased production of glycoproteins that are recognized by CTB (Fig. 3B). We therefore used MS to assess the N- and O-linked glycan composition on proteins isolated in triplicate from B3GALT5 KO or control cells by LC-MS/MS. Both N- and O-linked glycans extracted from B3GALT5 KO cells were larger than those from control cells (Fig. 4 and Supplementary Figs. 6A and 7A). Glycans from B3GALT5 KO cells were also more highly fucosylated. N-linked glycan structures with multiple fucose residues were detected (Supplementary Fig. 6B) and N-linked glycans with two or more fucose residues were enriched in B3GALT5 KO cells (Supplementary Fig. 6C). Even more dramatic differences were detected for O-GalNAc glycans. The mono-fucosylated, singly-extended core 2 O-glycan (Galβ(1-3/4)[Fucα1-2/3/4]GlcNAcβ(1-6)[Galβ(1-3)]GalNAcαSer/Thr) was the only fucosylated O-glycan detected in control cells whereas in B3GALT5 KO cells, an abundance of large, multiply-extended core 2 glycans displaying multiple fucose residues were detected (Fig. 4 and Supplementary Figs. 7A, 7B, and 7C). Thus, B3GALT5 KO cells are enriched in expression of fucosylated glycoproteins.

To assess the role of fucosylation in CT binding to B3GALT5 KO cells, we utilized CTB mutants that have a point mutation in either of the glycan binding pockets (Fig. 5A).^45^ The W88K point mutation in the canonical binding pocket abrogates CTB binding to the GM1 glycan but binding to fucosylated glycans is unaffected. Conversely, the H18L point mutation in the non-canonical binding pocket disrupts CTB binding to fucosylated glycans but binding to GM1 is unaffected. We analyzed binding of biotinylated CTB mutants alongside WT CTB to control and B3GALT5 KO cells by flow cytometry. In both B3GALT5 KO and control cells, the W88K CTB mutant retained almost all binding, whereas the H18L mutant exhibited significantly decreased binding relative to WT CTB (Fig. 5B). Ligand blot analysis confirmed that binding of CTB to both glycoproteins and glycolipids from control and B3GALT5 KO cells is dependent on the non-canonical pocket (Fig. 5C) whereas binding to GM1 depends on the canonical pocket (Supplementary Fig. 8A). To check the functional significance of the observed differences in CTB mutant binding, we employed the CTB-saporin internalization assay. Treatment of both control and B3GALT5 KO cells with W88K CTB-saporin resulted in little change in cell survival relative to WT CTB-saporin treatment (Supplementary Fig. 8B). Treatment of both control and B3GALT5 KO cells with H18L CTB-saporin resulted in a complete protection from saporin-induced cell death. Together, these data demonstrate that the non-canonical binding pocket is required for CTB binding to and internalization in both B3GALT5 KO and control cells but the canonical binding pocket is not.

### Fucosylation is required for cholera toxin intoxication of B3GALT5 KO cells

To further assess the functional significance of fucosylation in intoxication by CT, we used CRISPR/Cas9 to knockout the GDP-fucose transporter SLC35C1 in WT and B3GALT5 KO cells. In addition to being a top hit of our CRISPR screen analysis, SLC35C1 is necessary for fucosylation^34^. We confirmed that Le^x^ and other fucosylated glycans were absent from these cells (Supplementary Fig. 9A, 9B, 9C, and 9D). Using flow cytometry (Fig. 6A) and CTB lectin blot (Fig. 6B) analyses, we observed dramatically reduced CTB binding both SLC35C1 KO and B3GALT5+SLC35C1 dKO cells. To check the functional significance of the observed differences, we employed the CTB-saporin internalization assay. Both SLC35C1 and B3GALT5+SLC35C1 dKO cells were completed protected from CTB-saporin-induced cell death (Fig. 6C) but did not exhibit altered sensitivity to saporin alone (Supplementary Fig. 9E). Finally, to test if loss of CTB binding in SLC35C1 KO cells impacted intoxication, we treated cells with unlabeled CT and measured cAMP accumulation. We found that loss of SLC35C1 was protective against cholera intoxication even in the background of the sensitized B3GALT5 KO (Fig. 6D). Together, these results show that sensitization to CT in B3GALT5 KO cells is dependent on fucosylation.

**Figure 6.**
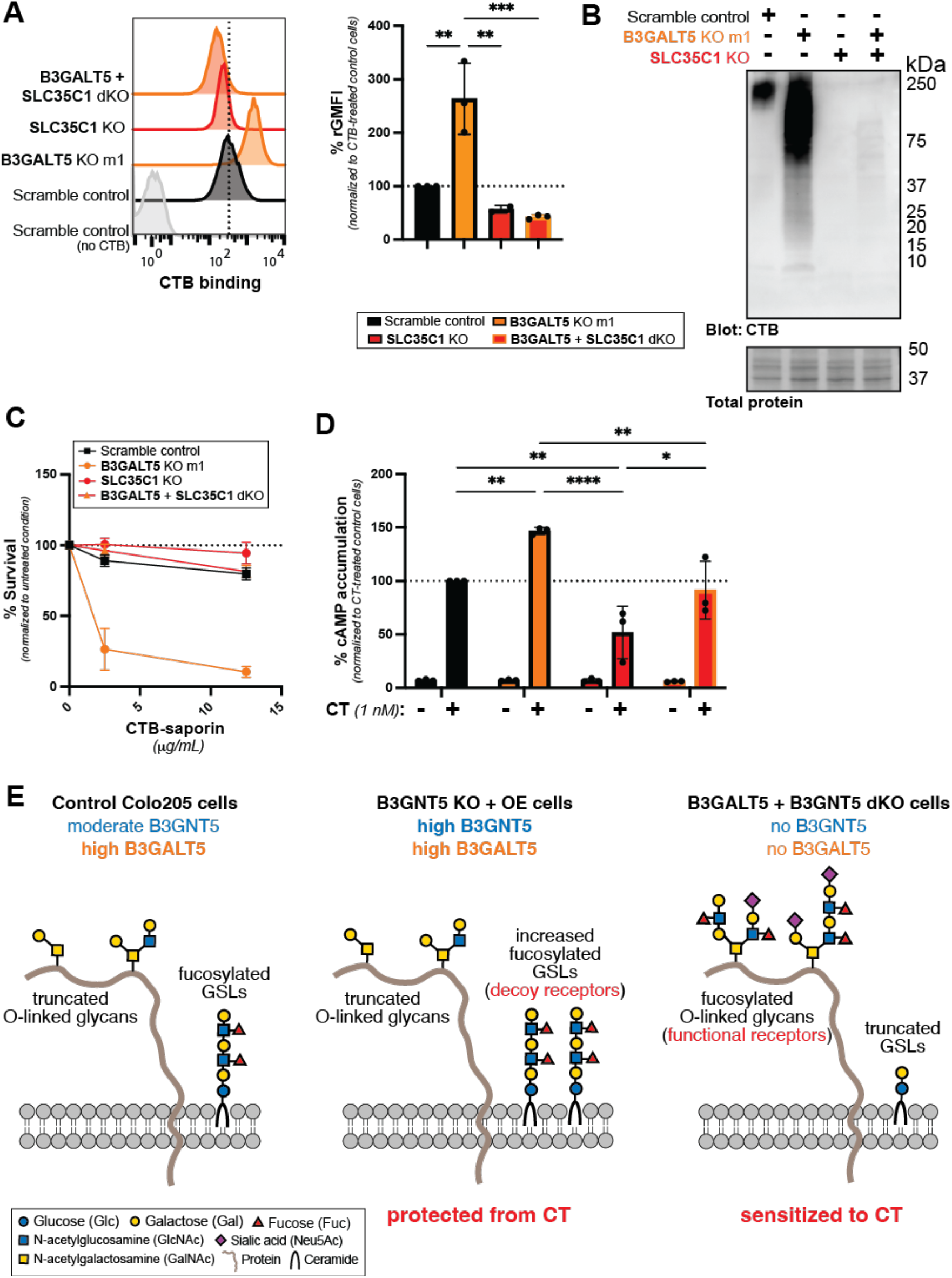
Fucosylated glycoproteins and fucosylated glycolipids play opposing roles in CT intoxication. (**A**) Representative histograms from the flow cytometry analyses of CTB (1 μg/mL) binding to surfaces of control, B3GALT5 KO m1, SLC35C1 KO, and B3GALT5+SLC35C1 dKO cells. Data shown are a single representative trial from 2-3 independent experiments. Quantification of gMFIs (bottom panel) from the 2-3 independent trials are normalized to the maximum signal in control cells. Error bars indicate mean ± SD. (**B**) Lysates from control, B3GALT5 KO, SLC35C1 KO, and B3GALT5+SLC35C1 double KO cells were analyzed by lectin blot, probing with CTB-biotin. Data shown are a single representative trial of 3 independent biological replicates. (**C**) Control, B3GALT5 KO m1, SLC35C1 KO and B3GALT5+SLC35C1 dKO cells were incubated for 72 h with increasing concentrations of CTB-saporin. Cell survival upon internalization of CTB-saporin was measured. Data shown are luminescence values normalized to the signal from the untreated condition for each cell type. Each datapoint is a biological replicate consisting of 3 averaged technical replicates. Error bars indicate mean ± SD of 2 biological replicates. (**D**) Control, B3GALT5 KO m1, SLC35C1 KO, and B3GALT5+SLC35C1 dKO cells were incubated for 1.5 h with CT (1 nM) after which accumulation of cAMP was measured. Data shown are inverse luminescence values normalized to total amount of cells then to the maximum signal in control cells. Each datapoint is a biological replicate consisting of 3 averaged technical replicates. Error bars indicate mean ± SD of 3 biological replicates. Statistical analyses for panel A performed by one-way ANOVA with Tukey correction and for panel D by two-way ANOVA for Tukey correction and * indicating p value between 0.01 and 0.05, ** indicating p value between 0.001 and 0.01, and *** indicating p value between 0.001 and 0.0001. (**E**) Model depicting how B3GNT5 and B3GALT5 regulate production of cholera toxin receptors. Control Colo205 cells express B3GNT5 and a high level of B3GALT5. These cells produce fucosylated GSLs and truncated O-linked glycans. Among the cell lines examined, B3GNT5 KO + OE cells were the most protected from CT. These cells express high levels of both B3GALT5 and B3GNT5. Due to high B3GALT5 expression, they do not produce fucosylated O-linked glycans. However, they produce abundant fucosylated GSLs due to high B3GNT5 expression. These fucosylated GSLs serve a protective role against CT intoxication. In contrast, B3GALT5 + B3GNT5 dKO cells were the most sensitized to CT. Due to the absence of B3GNT5 expression, fucosylated GSLs are not produced. However, the lack of B3GALT5 expression results in biosynthesis of fucosylated O-linked glycoproteins that promote CT intoxication. Representative glycan structures presented in this model are postulated based on O-linked glycomics analysis of control and B3GALT5 KO Colo205 cells and glycolipidomics analysis of control, B3GALT5 KO, and B3GNT5 KO Colo205 cells.

## DISCUSSION

Here, we report that the glycosyltransferases B3GALT5 and B3GNT5 are regulators of both CT binding and intoxication of Colo205 cells. These enzymes control the expression of fucosylated glycoconjugates that are recognized by CTB. B3GNT5 controls expression of fucosylated GSLs that act as decoy receptors, binding CTB and protecting cells from CT intoxication. Genetic disruption of B3GALT5 allows for biosynthesis of fucosylated O-linked glycoproteins that bind CTB and promote intoxication. The interplay of B3GALT5 and B3GNT5 expression determines which fucosylated structures are produced and whether they are displayed on glycoproteins versus GSLs.

Pioneering CRISPR inhibition and activation screens done by Gilbert and Horlbeck et al. used CT as a model system and also found activation of B3GNT5 to be protective against intoxication^46^. This consistency is especially striking since the cell line used, K562 lymphoblasts, expresses detectable levels of gangliosides including GM1.^47^ Indeed, the top glycosylation genes identified by Gilbert and Horlbeck et al. were involved in ganglioside biosynthesis (B3GALT4, B4GALNT1, and ST3GAL5). These genes were not enriched in our screen, suggesting that GM1 does not contribute significantly to CTB binding of Colo205 cells. Our data, however, do not exclude a role for GM1 or other non-fucosylated receptors in CT intoxication. For instance, we observe intoxication of SLC35C1 KO cells albeit at low levels, suggesting that other non-fucosylated receptors for CT might exist. GM1 was not detected in the MS analysis of GSLs extracted from Colo205 cells, which is consistent with minimal GM1 expression in normal human intestinal epithelial tissue.^6, 13^ Still, due to the high affinity of the interaction between GM1 and CTB, only small amounts may be necessary for intoxication. In addition, highly abundant receptors such as fucosylated glycoproteins could act in concert with low abundance receptors such as GM1 to drive intoxication.^27^ Indeed, even in the presence of detectable amounts of GM1, fucosylated glycoconjugates can affect CT activity ^31, 46^.

Future efforts will fill in additional molecular details regarding the fucosylated receptors recognized by CT. B3GALT5 KO cells exhibit increased fucosylation of both N- and O-linked glycans, however, our current data do not assess the relative importance of these two classes of glycans in CT intoxication. Furthermore, knockout of B3GALT5 results in a dramatic shift in O-glycan composition from mainly tumor-associated truncated core 1 O-glycans to diversely complex, fucosylated core 2 O-glycans. This change is associated with increased sensitivity to CT; however, normal human intestinal epithelia also display abundant core 3 extended O-glycan structures that were not examined here.^48, 49^ One limitation of the work presented here is that it was performed in a colorectal cancer cell line. However, a rstudy performed using human enteroids derived from jejunal biopsies also implicated fucose as playing a dual role in contributing to both functional and decoy receptors.^45^

Fucosylation at the intestinal epithelial cell surface has major implications in human health and disease ^50, 51^. Commensal bacteria induce fucosylation of the intestinal epithelia, which supports host-microbe symbiosis in the face of pathogenic challenge^52–55^. At the same time, certain pathogens have evolved to exploit host cell surface fucosylation. For instance, *Campylobacter jejuni* and *Salmonella typhimurium* use host fucose as an energy source^56–58^, and both bind to fucosylated blood group glycans to attach to the intestinal epithelium^59, 60^. Additionally, heat labile enterotoxin from *Escherichia coli* as well as norovirus and *Helicobacter pylori* exhibit a blood group dependence due to their ability to bind to and discriminate among different fucosylated glycans^61–76^. Together, these findings highlight the importance of understanding the regulation and composition of fucosylated structures at the human intestinal cell surface.

## METHODS

### Chemicals

Dimethyl sulfoxide (DMSO) was purchased from Sigma. (1R,2R)-1-phenyl-2-palmitoylamino-3-pyrrolidino-1-propanol (P4) was a gift from Prof. Ronald Schnaar (Johns Hopkins); stock concentrations were made at 5 mM in water. Bovine serum albumin (BSA) was purchased from Sigma. Skim milk powder was purchased from Fisher Scientific (catalog no. NC9121673). Carbohydrate-free blocking solution was purchased from Vector Laboratories (Burlingame, CA) (catalog no. SP-5040). COmplete ULTRA protease inhibitor tablets were purchased from Roche. Brefeldin A (≥ 99% pure) was purchased from Sigma (catalog no. B7651); stock concentrations were made at 5 mg/ml in DMSO. Forskolin (≥ 98% pure) was purchased from Sigma (catalog no. F6886); stock concentrations were made at 10 mM in DMSO. 1-Methyl-3-Isobutylxanthine (IBMX, ≥ 98% pure) was purchased from Cayman Chemicals (catalog no. 13347); stock concentrations were made at 50 mM in DMSO. Ro 20-1724 [4-(3-butoxy-4-methoxy-benzyl) imidazolidone] was purchased from Sigma Aldrich (catalog no. B8279); stock concentrations were made at 100 mM in DMSO.

### Endoglycoceramidase I (EGCase) expression and purification

*E.coli* BL21 (DE3) cells were transformed with a pET30 plasmid encoding *Rhodococcus triatomae* EGCase I (RhtrECI).^77^ A 1L culture of the transformed cells was induced with 0.1 mM IPTG at 16 °C for 18-20 h with continuous shaking at 200 rpm. Cells were harvested by centrifugation (8,980 *g* for 30 min) and resuspended in 40 mL lysis buffer (50 mM Tris pH 7.5 containing 300 mM NaCl and 0.1 % Triton X-100). Cells were disrupted by sonication for 5 min (60 amplitude with 10 seconds pulse on and 30 seconds pulse off). The supernatant was cleared by centrifugation (150,700 *g* at 4 °C for 1 h), then loaded on to 2 mL Ni-NTA column (Qiagen) pre-equilibrated with 10 column volumes of 50 mM Tris pH 7.5 containing 300 mM NaCl. EGCase I was eluted with 250 mM imidazole and dialyzed with TBS (50 mM Tris-HCl, pH 7.5 containing 150 mM NaCl). The homogeneity of purified EGCase I was analyzed by SDS-PAGE.

### Cell culture

The following reagents were used for general cell culture: fetal bovine serum (FBS) (catalog no. 16000), penicillin-streptomycin, TrypLE Express enzyme with phenol red (Gibco), Dulbecco’s Phosphate Buffered Saline (DPBS) (Sigma-Aldrich). Colo205 cells (ATCC) were maintained in RPMI-1640 media (Gibco) supplemented with 10 % FBS (v/v), and 1 % penicillin-streptomycin (Sigma). HEK293T/17 cells (ATCC) were maintained in Dulbecco’s Modified Eagle’s media (DMEM) supplemented with 20% FBS. Both cell lines were maintained at 37 °C, 5 % carbon dioxide in a water-saturated environment and were not used past passage number 45 or 15, respectively. The Countess automated cell counter (Life Technologies) was used for cell counting.

### Cholera toxin

Biotin-conjugated cholera toxin B subunit (CTB) used for the CRISPR screen was purchased from Life Technologies (catalog no. C-34779). Azide-free cholera holotoxin (CT) from *Vibrio cholerae* used for cAMP experiments was purchased from List Biological Laboratories (Campbell, CA; catalog no. 100B). Unlabeled CTB and CTB mutants used for flow cytometry experiments and biotin-conjugated CTB and CTB mutants used for internalization and ligand blot experiments were prepared as described.^45^

### Flow cytometry

Colo205 cells were seeded in a 10-cm tissue culture plate at a density of 2 x 10^5^ cells/mL and cultured before experiments for 48 h or for 72 h when seeded at a density of 1 x 10^5^ cells/mL. For experiments requiring inhibition of glucosylceramide glycosphingolipid biosynthesis, 1 x 10^5^ cells/mL were plated with the addition of P4 inhibitor at a final concentration of 1 μM and cultured for 72 h. Media containing floating cells was collected and 2 mL of 10 mM EDTA in DPBS was added to adherent Colo205 cells and incubated for 5 min at 37 °C. Adherent cells were then dislodged by gentle resuspension and then combined with previously harvested floating cell population. Cells were washed twice by resuspending in DPBS, centrifuged at 500 *g* for 3 min and EDTA removed by aspiration. The cells were resuspended in DPBS containing 0.1 % (w/v) BSA (DPBS/BSA) and 3.5 x 10^5^ cells were added per well to a V-bottom plate (Costar, catalog no. 3897). Cells were pelleted by centrifugation at 730 *g* for 5 min at 4 °C and washed twice by resuspension in 200 μL cold DPBS/BSA. Cells were incubated for 30 min on ice with 50 μL of 0, 0.0375, 0.075, 0.15, 0.3 or 0.6 μg/mL of unlabeled CTB (1 mg/mL) for dose responses. For all other CTB binding experiments, 50 μL of 1 μg/mL CTB was used. Cells were washed twice with 200 μL of cold DPBS/BSA and incubated for 30 min on ice with 50 μL of 1:500 dilution of anti-beta subunit cholera toxin (Abcam, catalog no. ab 34992). For antibody staining, 50 μL of 1:250 mouse anti-human CD15 clone HI98 [anti-Lewis x] (BD Biosciences, catalog no. 555400BD) or mouse-anti-human blood group Lewis a clone 7LE (Abcam, catalog no. ab3967). For staining with fucose-binding protein *Aleuria aurantia* lectin (AAL), 50 μL of 1:500 biotinylated AAL (Vector Labs) was used. The cells were washed twice, then stained with propidium iodide (PI) (Sigma-Aldrich, catalog no. P4170) at a final concentration of 1 μg/mL along with 50 μL of 1:500 dilution of respective secondary reagents for 30 min on ice. For CTB, the secondary reagent was donkey anti-rabbit IgG Alexa Fluor® 647 (Abcam, ab150075); for Le^x^, the secondary antibody was goat anti-mouse IgM Alexa Fluor® 647 (Abcam, ab150123); for Le^a^, the secondary antibody was goat anti-mouse IgG Alexa Fluor® 633 (Invitrogen, A-21050); for AAL, the secondary reagent was streptavidin-APC (Thermo Fisher, S868). After two washes, cells were analyzed on a FACSCalibur flow cytometer (BD Biosciences, UT Southwestern Flow Cytometry Core facility). Dead cells were excluded based on PI staining on the FL3 channel and the fluorescence intensity of the live population was determined on the FL4 emission channel.

### CRISPR screen

Human Brunello genome-wide CRISPR KO pooled library (gift from David Root and John Doench, Addgene catalog no. 73178) was amplified by electroporation into Stbl4 electrocompetent cells (Thermo Fisher Scientific, catalog no. 11635-018). Briefly, 400 ng of pooled plasmid library was added to 100 uL Stbl4 cells. 25 μL of cells were added to an electroporation cuvette (0.1 cm gap, Bio-Rad, catalog no. 165-2089), and electroporated using the Ec1 setting (1.8 kV) of the MicroPulser from Bio-Rad (catalog no. 1652100). 1 mL pre-warmed Super Optimal broth with Catabolite repression media (SOC, Thermo Fisher Scientific, catalog no. 15544034) was immediately added to cells, which were then transferred to round-bottom 14 mL tube (Falcon, catalog no. 352059). This process was repeated three times with the remaining Stbl4 cells. SOC was added to the combined electroporated cells to a final volume of 10 mL, which was split equally into two round-bottom 14 mL tubes and shaken for 1 hour at 30 °C. 2.5 mL of cells were spread evenly on each of four 500 cm^2^ bioassay agar/carbenicillin (100 ug/ml) plates and grown at 30 °C for 16 h. Colonies were scraped and plasmid DNA was prepared using the HiSpeed Plasmid Maxi kit (Qiagen, catalog no. 12662) with the following modifications: buffers P1, P2 and P3 were added directly to the 50 ml conical tube and centrifuged to pellet lysed debris before plunger was added, and DNA was eluted with buffer pre-warmed to 50 °C.

To generate lentivirus, 7.5 x 10^6^ low passage HEK293T/17 cells were plated on each of ten 15-cm dishes in 30 mL complete media. After 24 h, 48 uL TransIT-293 transfection reagent (Mirus Bio LLC, catalog no. MIR 2704) diluted with 1300 uL serum-free media (SFM) was incubated for 5 min at room temperature and then combined with a mix of 8 µg purified plasmid library DNA, 8 µg psPAX2 packaging plasmid DNA and 1 µg pMD2.G envelope plasmid DNA (both gifts from Didier Trono, Addgene plasmid no. 12260 and 12259, respectively). After incubating for 30 min, the mix was evenly split and added dropwise to each plate of HEK293T/17 cells. Viral supernatant was collected 72 h post infection, filtered through a 0.4 μm filter and then titrated in 6 well plates containing 2 x 10^6^ Colo205 cells to determine the amount of virus needed to infect cells at an MOI of 0.3. Briefly, cells were resuspended in media with 8 µg/ml polybrene and increasing volumes of lentivirus (0, 50, 100, 200, 500, or 1000 uL brought up to 2 mL with complete media). Cells were then centrifuged at 1000 *g* for 2 h at 33 °C and then transferred to an incubator and maintained at 37 °C in 5 % carbon dioxide for 16 h. Each well (containing ∼ 4 x 10^6^ cells) was then harvested and 2.5 x 10^4^ cells were plated in triplicate in wells of a white 96-well plate with a clear bottom (Costar Laboratories, Cambridge, MA). 24 h after passaging, library-expressing cells were selected with 5 µg/mL puromycin. 48 h post-selection, cell death was measured using the CellTiter-Glo 2.0 luminescent cell viability assay kit (Promega), and the virus volume condition (100 uL) resulting in ∼30% viability was selected for use.

2 x 10^6^ Colo205 cells were spinfected in each well of eight 6-well plates (9.6 x 10^7^ cells total to ensure 300x coverage of the pooled library post-selection) with 100 uL of virus brought up to 2 mL with complete media containing 8 µg/mL polybrene, as determined by titration described above. 24 h post infection, cells were harvested, resuspended in fresh media and expanded in twenty 15-cm dishes. 24 h after passaging, library-expressing cells were selected with 5 µg/mL puromycin. Cells were harvested 0 and 14 days post puromycin selection, consistently maintaining 300x library coverage. After harvesting on day 14, cells were centrifugated at 500*g* for 3 min at 4 °C and washed twice in DPBS. An input control population was set aside, and the remaining cells were resuspended in DPBS containing 2 % FBS (FACS buffer). 7 x 10^6^ cells were added to each of four round-bottom polystyrene 12 x 75 mm tubes (Corning, catalog no. 352008) and then washed twice by resuspension in 4 mL cold FACS buffer. Cells were then incubated for 30 min on ice with 1 mL of biotin-CTB (40 µg/mL). After two washes, cells were incubated with 1 mL of a 7.7 μg/mL dilution of fluorescein (DTAF) streptavidin (Jackson Immuno Research, West Grove, PA) (catalog no. 016-010-084). After a final two washes, cells were sorted and the top and bottom 1 % fluorescent cells were collected using the FACSAria flow cytometer (BD Biosciences, UT Southwestern Flow Cytometry Core facility).

Immediately after sorting, both the input and sorted cell populations were pelleted and genomic DNA (gDNA) was isolated from each using the Blood and Cell Culture DNA Maxi Kit (Qiagen, catalog no. 13362) according to the manufacturer’s instructions. To amplify sgRNA sequences for Next Generation Sequencing (NGS), four parallel 100 μL PCR reactions were initially run for each condition, and pooled. Each 100 μL PCR reaction contained 1 - 2 μg of gDNA, Ex Taq polymerase (Takara Bio, catalog no. RR01CM) and the PCR1 primer pair listed below. Amplified DNA was then indexed and barcoded for NGS in another PCR reaction including 5 μL of DNA from the initial PCR, Ex Taq polymerase, pooled P5 and barcoded P7 primers, as previously described^78^. DNA was then pooled and purified for sequencing using AMPure XP beads (Agencourt). 300 μL of pooled PCR product was mixed with 150 μL beads and incubated for 5 minutes to remove gDNA. Beads bound to gDNA were pelleted on the DynaMag-2 Magnet (Invitrogen) and the supernatant was then mixed with 540 μL of fresh beads and incubated for 5 minutes to bind the PCR products. After discarding the supernatant, beads were washed twice with 1 mL 70 % ethanol and then dried for approximately 5 min. Bound DNA was eluted from the beads using 300 μL sterile, nuclease-free water. Samples were sequenced using the Illumina NextSeq 500 with the read configuration as single end 75bp at UT Southwestern McDermott NGS core. Data were analyzed with MAGeCK.^79^ The gene set enrichment analyses of the identified genes were performed with WebGestalt.^80^ The Manhattan dot plots of the screen results were drawn with R package qqman (v. 0.1.8).^81^

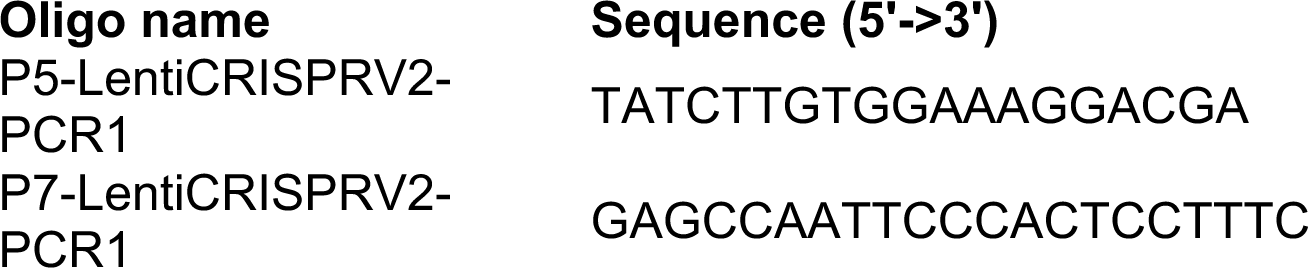

### Construction of cell lines

Overexpression plasmids for the rescue of B3GALT5 and B3GNT5 KOs were generated by Gibson cloning using the primers listed below designed using the NEBuilder software from New England Biolabs (NEB) to amplify the open reading frame (ORF) of each gene (Origene, catalog no. RC211183L3 and RC206965L3, respectively) while introducing a single bp change to the PAM sequence that does not result in a change to amino acid sequence. Amplification was done using the Q5 High-Fidelity 2X Master Mix (NEB, catalog no. M0492S) according to manufacturer’s instructions. Amplified ORFs were then purified using the MinElute PCR Purification Kit (Qiagen, catalog no. 28004) according to the manufacturer’s instructions. Purified PCR products were ligated using the HiFi DNA Assembly master mix (New England Biolabs, catalog no. E2621S) into a Blasticidin-resistance encoding lentiviral vector (Addgene, plasmid no. 52962) cut with BamHI-HF and AgeFI (NEB, catalog no. R3136S and R0652S, respectively) according to manufacturer’s guidelines. Plasmids were transformed into DH5α competent cells according to manufacturer’s instructions, which were expanded for maxiprep using the ZymoPure II maxiprep kit (Zymo Research, catalog no. D4204). Lentivirus was prepared as described for the CRISPR screen, but scaled for production in individual 10-cm dishes.

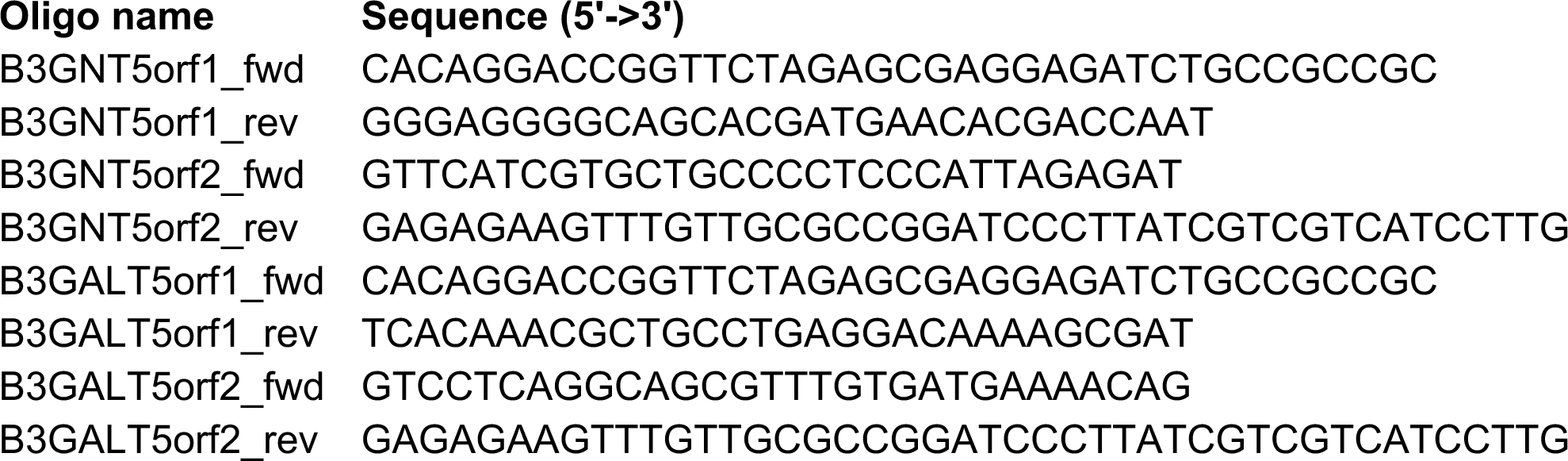

All-in-one CRISPR KO plasmid lentivirus for the expression of Cas9 as well as predesigned sgRNAs targeting B3GALT5, B3GNT5, SLC35C1, or a non-targeting scramble control sgRNA (Sigma, catalog no. HSPD0000061666, HSPD0000119328, HSPD0000095207, or NegativeControl1, respectively) were used to spinfect Colo205 cells as described above. Monoclonal populations were obtained using FACS in a sterile environment (UT Southwestern Flow Cytometry Core) to collect single, GFP-positive cells in each well of a 96-well tissue culture plate containing 200 μL complete media supplemented with extra FBS (20 % v/v total). Double KO (dKO) cells were generated in the same way in B3GALT5 single KO cells. Monoclonal overexpression cell lines were isolated by limiting dilution.

To sequence the CRISPR-targeted genes, gDNA was isolated from 5 x 10^6^ Colo205 cells using the Blood and Cell Culture DNA Maxi Kit (Qiagen, catalog no. 13362) according to the manufacturer’s instructions. The respective gene targets were amplified by polymerase chain reaction using Ex Taq polymerase (Takara Bio, catalog no. RR01CM) according to manufacturer’s instructions. The forward and reverse primers for each sequenced gene are listed below. The PCR products were purified using the NucleoSpin Gel and PCR Clean-up Mini Kit (Macherey-Nagel, catalog no. 740609.50) according to the manufacturer’s instructions. Purified DNA was ligated into a pGEM-T Easy pre-linearized vector (Promega, catalog no. A1360) and transformed into DH5α competent cells. Plasmids were extracted from single colonies and then submitted to the UT Southwestern Sanger Sequencing Core. Sequence data are presented in Supplementary Table 1. For all KO cell lines, except the B3GALT5 + SLC35C1 dKO, two different indels were identified, each in duplicate or triplicate. For KO of SLC35C1 in the B3GALT5 KO line, four separate clones harbored the same SLC35C1 indel. Loss of SLC35C1 activity was inferred by the complete loss of anti-Le^x^ and AAL binding to these cells.

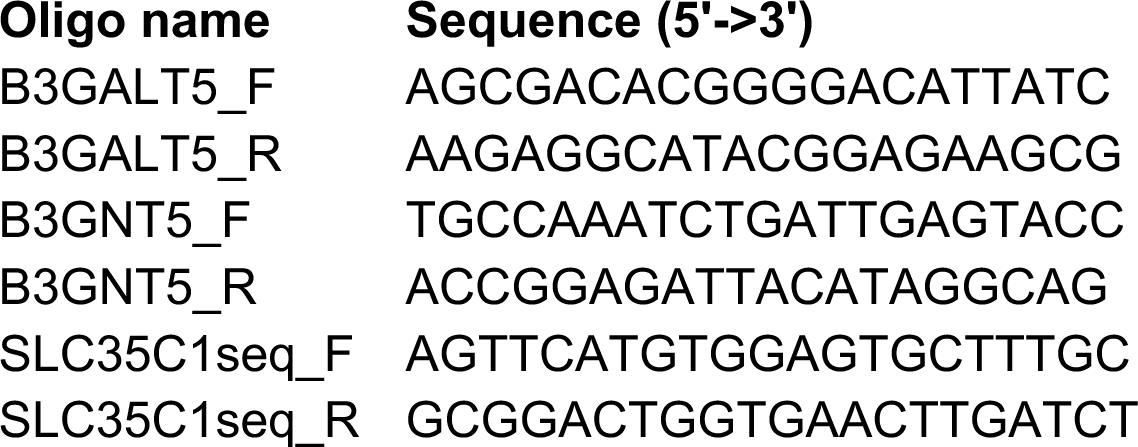

### Measurement *of CTB internalization*

2.5 x 10^3^ Colo205 cells in 20 µL of serum-free media were plated in each well of a white-walled, 96-well clear-bottom plate (Costar Laboratories, Cambridge, MA). Cells were cultured for 72 h in serum-free media with 20 µL of CTB-saporin or saporin alone (0, 1.25, 2.5 and 12.5 µg/mL final concentrations) in triplicate at 5 % CO_2_ at 37 °C. CTB-saporin complexes were prepared according to manufacturer’s instructions. Briefly, 12.5 µL of 1 mg/mL biotinylated CTB was combined with 2.5 µL of streptavidin-saporin (Advanced Targeting Systems, IT-27) and then diluted with serum-free media to a volume of 50 µL to generate a 25 µg/mL CTB-saporin solution. After 72 h, cell death was measured using the CellTiter-Glo 2.0 (CTG) luminescent cell viability assay kit (Promega).

### Measurement of cAMP

2.5 x 10^4^ Colo205 cells in 20 µL were plated in each well of a white 96-well plate with clear bottom (Costar Laboratories, Cambridge, Mass). Triplicate wells were incubated with 10 µL Brefeldin A (1 µg/ml) in serum-free media supplemented with IBMX (500 µM) and Ro-20-1764 (100 µM) (induction buffer). All other wells were treated with 10 µL vehicle control. After 30 min, BFA- or vehicle control-treated cells were treated with 10 µL cholera holotoxin (CT) diluted in induction buffer (0 or 2 nM) in triplicate at 5 % CO_2_ at 37 °C for 90 min. Forskolin diluted in induction buffer (10 µM) was added to vehicle control-treated wells in triplicate and incubated at 5 % CO_2_ at 37 °C for 15 min prior to measurement of cAMP accumulation. Accumulated cAMP was measured using the cAMP-glo Max Assay Kit (Promega), according to manufacturer’s instructions with modification to incubation times (30 min incubation with cAMP-glo ONE buffer and 20 min incubation with the kinase-Glo buffer). Luminescence values were normalized by amount of live cells in vehicle control-treated wells (CellTiter-Glo 2.0 Assay Kit, Promega). CTG reagent was added to vehicle control-treated wells in triplicate 15 min after addition of the cAMP-glo Max Assay kit ONE buffer.

### Lectin blot and immunoblot analyses

Colo205 cells (4 x 10^6^) were lysed in 200 μL RIPA buffer (50 mM Tris-HCl pH 8; 150 mM NaCl; 0.01 % (v/v) SDS; 0.5 % (v/v) sodium deoxycholate; 1 % (v/v) IGEPAL CA-630; 1X protease inhibitor). Protein content was quantified with a BCA assay kit (Thermo Fisher Scientific) against a BSA standard curve for normalization. For protease digestions, 20 μg of lysate was incubated with 500 μg/mL proteinase K in RIPA buffer for 48 h at 37 °C. EGCase digestions were performed by incubating 20 μg of lysate with 40 μg of enzyme in 50 mM sodium acetate buffer, pH 5.2 supplemented with 0.2 % Triton X-100 overnight at 37 °C. For fucosidase treatment, 20 μg of lysate was incubated with 1 μL of 100 U/μL FucosExo (Genovis, catalog no. G1-FM1-020) for 2 h at 37 °C. 20 μg of lysate was then denatured in 1X SDS loading dye (250 mM Tris-HCl, pH 6.8; 0.08 % (w/v) SDS; 0.0004 % (w/v) bromophenol blue; 40 % (v/v) glycerol; 5 % (v/v) 2-mercaptoethanol) for 10 min at 95 °C for all conditions. The samples were resolved on a 4-20 % stain-free gradient SDS-PAGE gel (Bio-Rad Laboratories) and activated for 1 min using the ChemiDoc MP Imaging system (Bio-Rad Laboratories). Activated protein was transferred to a PVDF membrane (ED Millipore) using the Trans-Blot SD Semi-Dry Transfer Cell (Bio-Rad Laboratories), according to the manufacturer’s guidelines at 15 V for 45 min and total protein stain was imaged using the ChemiDoc MP. After blocking at room temperature for 1 h, membranes were probed overnight at 4 °C with 1:2500 dilution of biotinylated CTB; 1:1000 dilution of mouse anti-human CD15 clone HI98 [anti-Lewis x] (BD Biosciences, catalog no. 555400BD), or biotinylated AAL lectin (Vector Laboratories). Membranes probed with CTB and anti-Lewis x were blocked and incubated in 5 % non-fat milk prepared in 1X TBS containing 0.05 % (v/v) Tween-20 (TBS-T). When probing with AAL, membranes were blocked and incubated with 1X Carbohydrate-free solution prepared in water. Specific binding of lectins and Lewis x antibody was assessed after incubating membranes with HRP-conjugated streptavidin or goat anti-mouse IgM secondary antibody (1:5000 dilution), respectively. HRP-signal was detected using the SuperSignal West femto Chemiluminescent substrate diluted in SuperSignal West Pico PLUS Chemiluminescent substrate (Thermo Fisher Scientific) and ChemiDoc MP Imaging system (Bio-Rad Laboratories).

### Glycosphingolipid Isolation

Preparation of glycolipids from whole cells was carried out using previously described methods.^82^ Pellets of cells were disrupted with probe sonication between extraction of the total lipids with 4:8:3 Chloroform:Methanol:Water (CMW). The mixture was bath-sonicated in ice before centrifugation to pellet the proteinaceous material to the bottom. The supernatant was then removed to a clean glass tube. Fresh aliquots of CMW were added 2 more times with sonication, each time removing the supernatant. The combined supernatant, which contains the crude lipid extraction, was then dried down, then reconstituted in 0.5 M NaOH in Methanol (MeOH:Water 95:5 v/v), sonicated to redissolve the lipids, then incubated at 37 °C overnight to saponify the phosphoglycerolipids. The next day an equal volume of 5 % acetic acid was added to the reaction to neutralize and halt saponification, and the sample was desalted on a tC18 SPE cartridge (Waters, Milford, MA), resulting in the isolated glycosphingolipid (GSL) material. Free fatty acids were removed from the sample by drying the GSL material by addition and removal of hexane. The sample was thoroughly dried prior to further analysis.

### N- and O-Glycan release

Cell pellets were resuspended in ammonium bicarbonate and passed several times through a 23-gauge syringe to homogenize. Cells were further lysed with probe sonication and the lysate was denatured by adding 10 µL of denaturing buffer (NEB PNGase F kit, P0709) and heating at 100 °C for 5 min. Salts were removed with a pre-washed 10 kDa MWCO filter and exchanged into 50 mM ammonium bicarbonate. Whole proteins were removed from the top of the MWCO filter and resuspended in 50 mM ammonium bicarbonate. N-glycans were released with PNGase F (NEB, P0709) for 20 h at 37 °C and the released N-glycans were isolated by passage through a pre-washed 10 kDa MWCO filter. The remaining protein including O-linked glycans remained in the top of the filter and were removed before treated with 1 M sodium borohydride in 50 mM sodium hydroxide. O-linked glycans were released with beta-elimination at 45 °C overnight. The basic solution was neutralized with a 10 % acetic acid solution, desalted through a packed Dowex column (50 W x8-100, Sigma Aldrich, St. Louis, MO) then lyophilized. Borates were removed from the sample with repeated additions of 9:1 methanol: acetic acid under a nitrogen flow. The material was then resolubilized and pass through a C18 SPE cartridge to further purify the material before permethylation and analysis.

### Permethylation and Mass Spectrometry (MS) Analysis

The GSL material was permethylated using previously described methods.^83–85^ Briefly, dried samples were reconstituted in DMSO and treated with a mixture of methyl iodide in a DMSO/NaOH slurry. After quenching the reaction with the addition of water, the permethylated analytes were extracted out of the solution by addition of dichloromethane.

MS analysis was carried out by direct infusion MS or LC-MS. Permethylated glycans and glycolipids were reconstituted in a 50:50 MeOH:H_2_O mixture containing 1 mM NaOH which acts as a charge carrier. The solution was analyzed on a Thermo Orbitrap Fusion Tribrid Mass Spectrometer with direct infusion at a flow rate of 1-2 µL/min and analyzed in positive ion mode. An automated method that collects a full MS spectra at 120 000 resolution followed by data dependent MS/MS fragmentations (CID) at 60 000 resolution was used to collect data for the glycosphingolipids. Data for the permethylated N- and O-glycans were collected by LC-MS on a Thermo Orbitrap Fusion Tribrid Mass Spectrometer in line with an Ultimate 3000RSLCnano LC-system. A commercial C18 column (Thermo Fisher; Cat 164568) of 15-cm x 75µm id filled with 3-µm C18 material) was used for analysis. Injections of permethylated N- or O-linked glycans were carried out from low to high acetonitrile in water with 1 mM sodium acetate. The precursor ion scan was acquired at 120 000 resolution in the Orbitrap analyzer and precursors at 3 sec were selected for MS/MS fragmentation (CID fragmentation) in the Orbitrap at 30 000 resolution.

### Statistical analysis

Statistical analyses were performed using Prism 9 software (GraphPad). * indicates p value between 0.01 and 0.05, ** 0.001 and 0.01, *** 0.001 and 0.0001, and **** <0.0001.

## ACKNOWLEDGEMENTS

We thank Dr. Akshi Singla, Dr. Débora Andrade Silva, and Mary Burns for comments on the manuscript. We thanks Emanuela Capota and Dr. Débora Andrade Silva for technical support, and Dr. Haaris Khan and Prof. Michael Shiloh for advice and reagents. We acknowledge support from the NIH (R01GM090271 and R35GM145599 to JJK and NIH R24GM137782 to PA) and the Welch Foundation (I-1686 to JJK). We thank Christopher Cairo (University of Alberta) for the RhtrECI plasmid and Ronald Schnaar (Johns Hopkins) for the P4 inhibitor.

