## Supplementary Figures 1 - 9 and Supplementary Table 1 for "Fucosylated glycoproteins and fucosylated glycolipids play opposing roles in cholera intoxication"

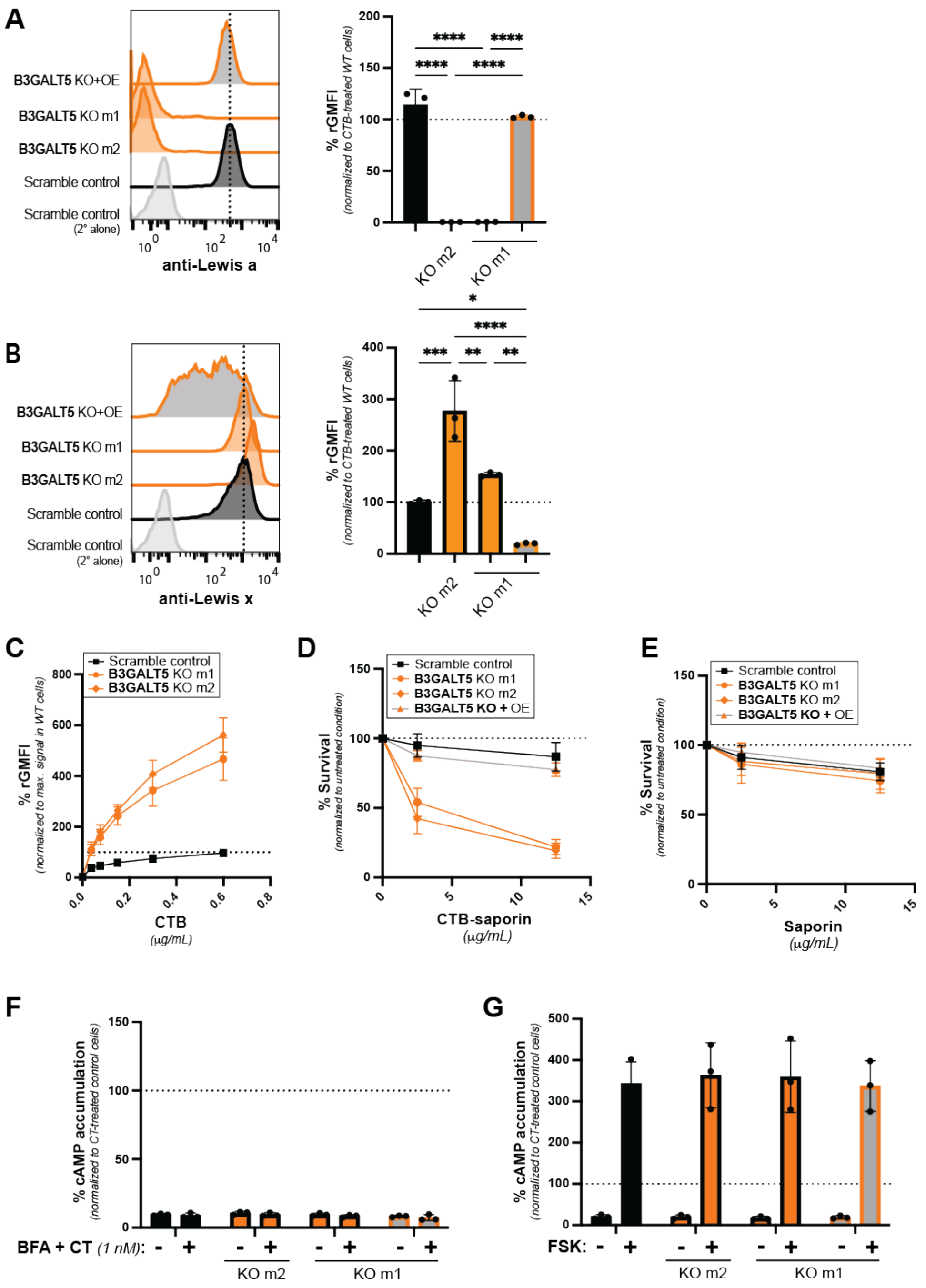


**Supplementary Figure 1. Characterization of B3GALT5 KO and B3GALT5 KO + OE cells**

Representative histograms from the flow cytometry analyses of cell surface binding of Lewis a antibody (**A**) or Lewis x antibody (**B**) to control, B3GALT5 KO m1, m2 and KO+OE cells. Bar graphs show quantification from 3 independent trials. (**C**) Quantification of gMFI from flow cytometry analyses of cells treated with increasing concentrations of CTB. Data shown are from 3 independent trials and normalized to the maximum APC signal in WT cells. Error bars indicate mean ± SD. Control, B3GALT5 KO m1 and KO+OE cells were incubated for 72 h with increasing concentrations of CTB-Saporin (**D**) or unconjugated saporin (**E**). Cell survival upon internalization of CTB-saporin measured using the Cell Titer-Glo 2.0 assay. Data shown are luminescence values normalized to the signal from the untreated condition for each cell type. Each datapoint is a biological replicate consisting of 3 averaged technical replicates. Error bars indicate mean ± SD of 3 biological replicates. (**F**) Cells pretreated with brefeldin A (BFA) or a vehicle control for 0.5 h were incubated for 1.5 h with CT (1 nM) or buffer alone. Accumulation of cAMP was measured. Data shown are inverse of luminescence values normalized first to the total amount of cells plated for each cell line, then to the signal in CT-treated control cells. Each datapoint is a biological replicate consisting of 3 averaged technical replicates. Error bars indicate mean ± SD of 3 biological replicates. (**G**) Control, B3GALT5 KO m1, m2 and KO+OE cells were treated with forskolin (10 µM) for 0.5 h and then analyzed as in panel F. Statistical analyses for panels A and B by one-way ANOVA with Tukey correction and for panels F and G by two-way ANOVA with Tukey correction (* indicates p value between 0.01 and 0.05, ** 0.001 and 0.01, *** 0.001 and 0.0001, and **** ≤0.0001).

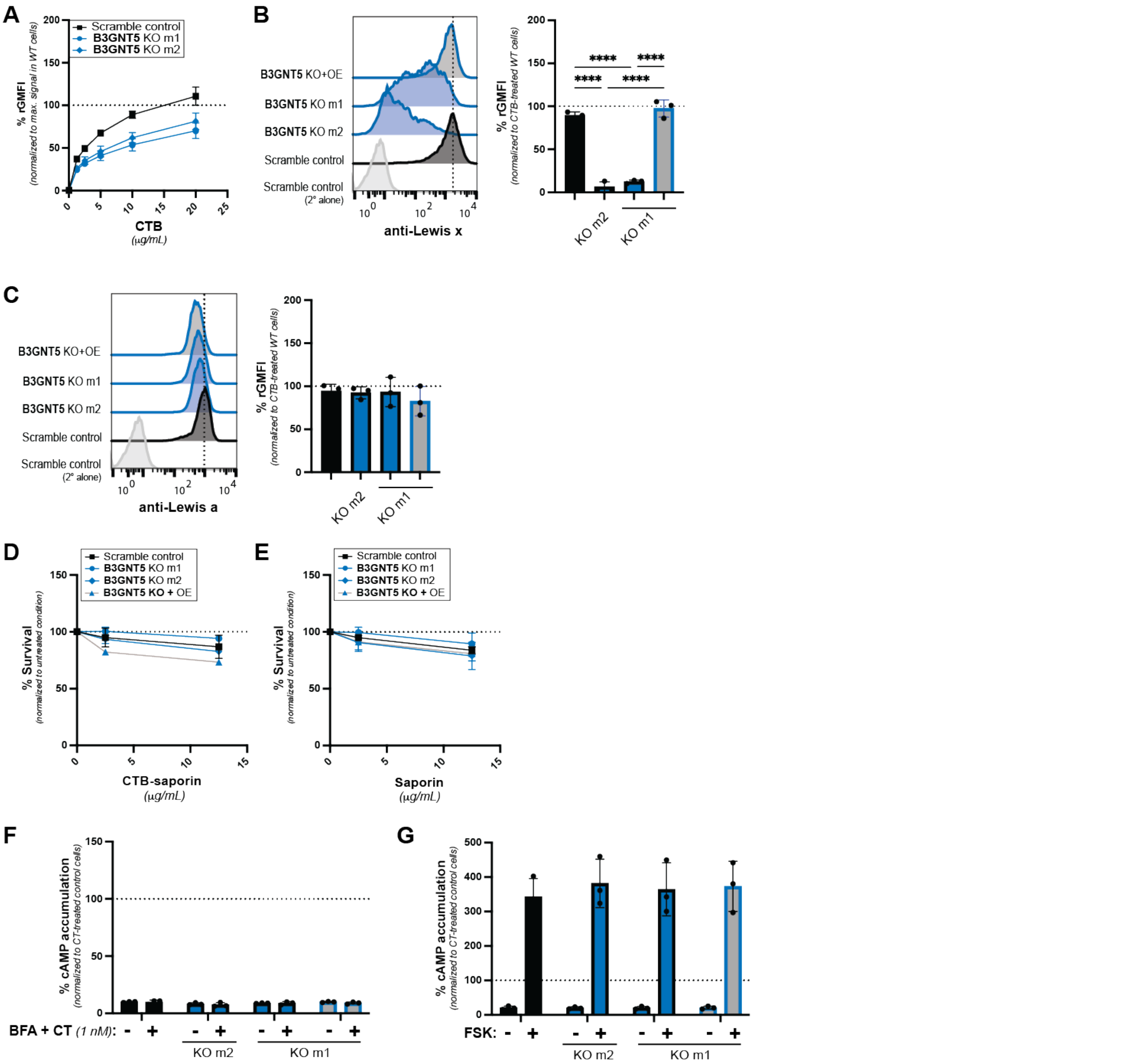


**Supplementary Figure 2. Characterization of B3GNT5 KO and B3GNT5 KO + OE cells**
(**A**) Quantification of gMFI from flow cytometry analyses of cells treated with increasing concentrations of CTB. Data shown are from 3 independent trials and normalized to the maximum APC signal in WT cells. Error bars indicate mean ± SD. Representative histograms from the flow cytometry analyses of cell surface binding of Lewis x antibody (**B**) or Lewis a antibody (**C**) to control, B3GNT5 KO m1, m2 and KO+OE cells. Bar graphs show quantification from 3 independent trials. Control, B3GALT5 KO m1 and KO+OE cells were incubated for 72 h with increasing concentrations of CTB-Saporin (**D**) or unconjugated saporin (**E**). Cell survival upon internalization of CTB-saporin measured using the Cell Titer-Glo 2.0 assay. Data shown are luminescence values normalized to the signal from the untreated condition for each cell type. Each datapoint is a biological replicate consisting of 3 averaged technical replicates. Error bars indicate mean ± SD of 3 biological replicates. (**F**) Cells pretreated with brefeldin A (BFA) or a vehicle control for 0.5 h were incubated for 1.5 h with CT (1 nM) or buffer alone. Accumulation of cAMP was measured. Data shown are inverse of luminescence values normalized first to the total amount of cells plated for each cell line, then to the signal in CT-treated control cells. Each datapoint is a biological replicate consisting of 3 averaged technical replicates. Error bars indicate mean ± SD of 3 biological replicates. (**G**) Control, B3GALT5 KO m1, m2 and KO+OE cells were treated with forskolin (10 µM) for 0.5 h and then analyzed as in panel F. Statistical analyses for panels B and C by one-way ANOVA with Tukey correction and for panels F and G by two-way ANOVA with Tukey correction (* indicates p value between 0.01 and 0.05, ** 0.001 and 0.01, *** 0.001 and 0.0001, and **** ≤0.0001).

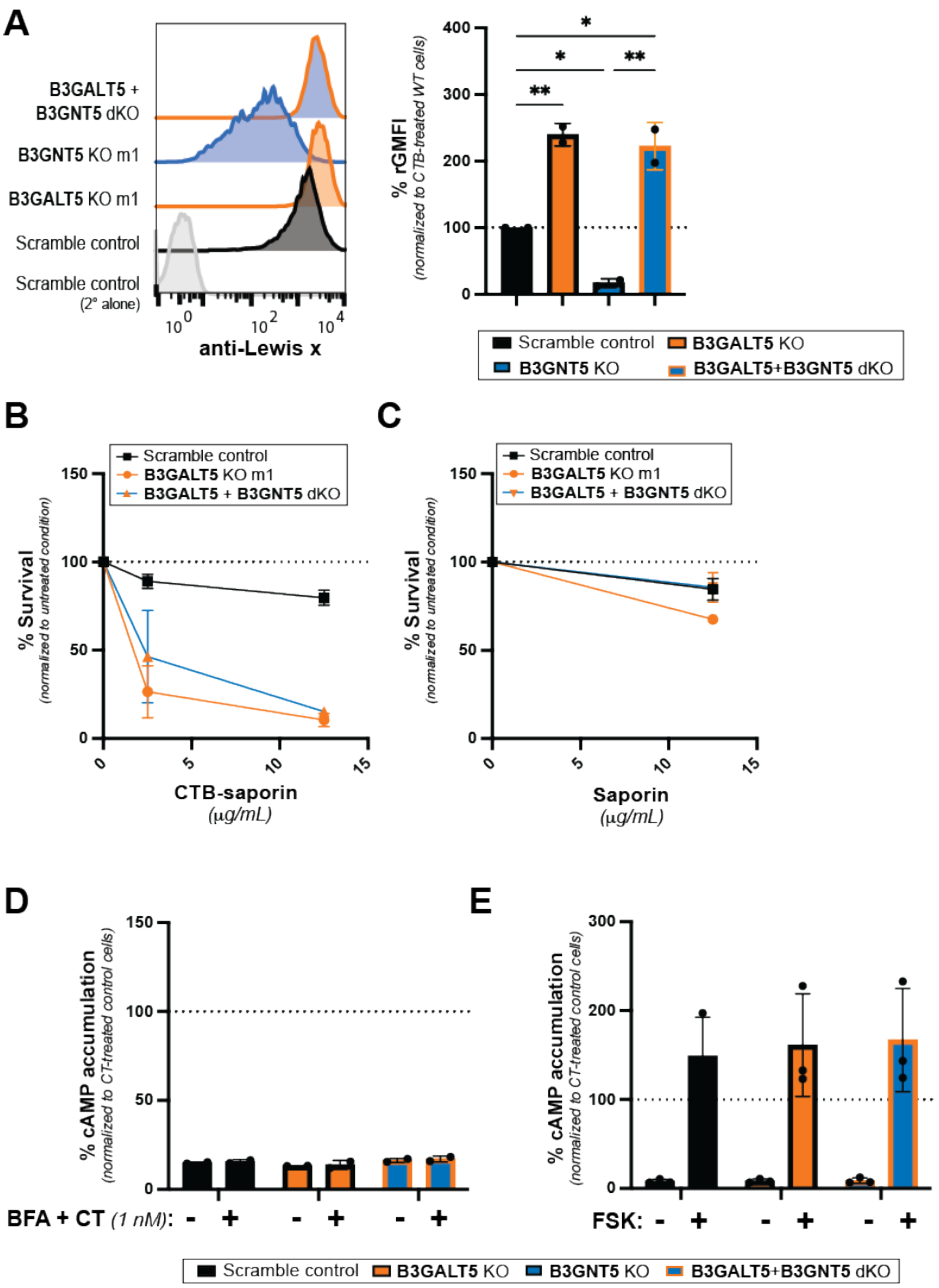


**Supplementary Figure 3. Characterization of B3GALT5 + B3GNT5 dKO cells.**
(**A**) Representative histograms from the flow cytometry analyses of cell surface binding of Lewis x antibody to control, B3GALT5 KO, B3GNT5 KO, and B3GALNT5 + B3GNT5 dKO cells. Bar graphs show quantification from 3 independent trials. Control, B3GALT5 KO, and B3GALNT5 + B3GNT5 dKO cells were incubated for 72 h with increasing concentrations of CTB-Saporin (**B**) or unconjugated saporin (**C**). Cell survival upon internalization of CTB-saporin measured using the Cell Titer-Glo 2.0 assay. Data shown are luminescence values normalized to the signal from the untreated condition for each cell type. Each datapoint is a biological replicate consisting of 3 averaged technical replicates. Error bars indicate mean ± SD of 3 biological replicates. (**D**) Cells pretreated with brefeldin A (BFA) or a vehicle control for 0.5 h were incubated for 1.5 h with CT (1 nM) or buffer alone. Accumulation of cAMP was measured. Data shown are inverse of luminescence values normalized first to the total amount of cells plated for each cell line, then to the signal in CT-treated control cells. Each datapoint is a biological replicate consisting of 3 averaged technical replicates. Error bars indicate mean ± SD of 3 biological replicates. (**E**) Control, B3GALT5 KO m1, m2 and KO+OE cells were treated with forskolin (10 µM) for 0.5 h and then analyzed as in panel D. Statistical analyses for panel A by one-way ANOVA with Tukey correction and for panels D and E by two-way ANOVA with Tukey correction (* indicates p value between 0.01 and 0.05, ** 0.001 and 0.01, *** 0.001 and 0.0001, and **** ≤0.0001).

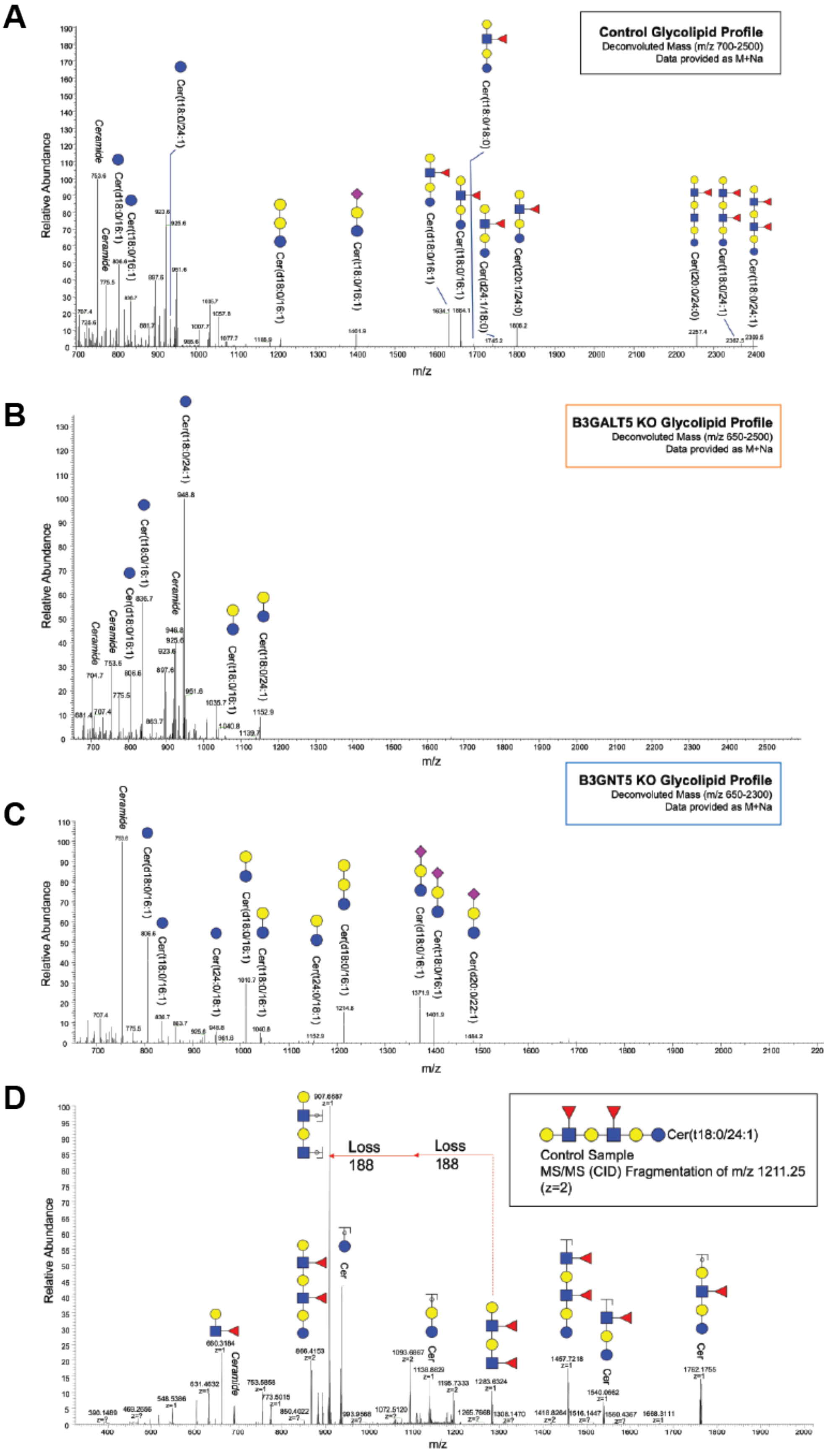


**Supplementary Figure 4. Fucosylated lacto-series GSLs detected in control but not KO cell lines.**
MS analysis of GSLs from control (**A**), B3GALT5 KO (**B**), and B3GNT5 KO (**C**) cells. (**D**) Example MS/MS spectrum of a fucosylated lacto-series glycan detected in control cells, confirming structure.

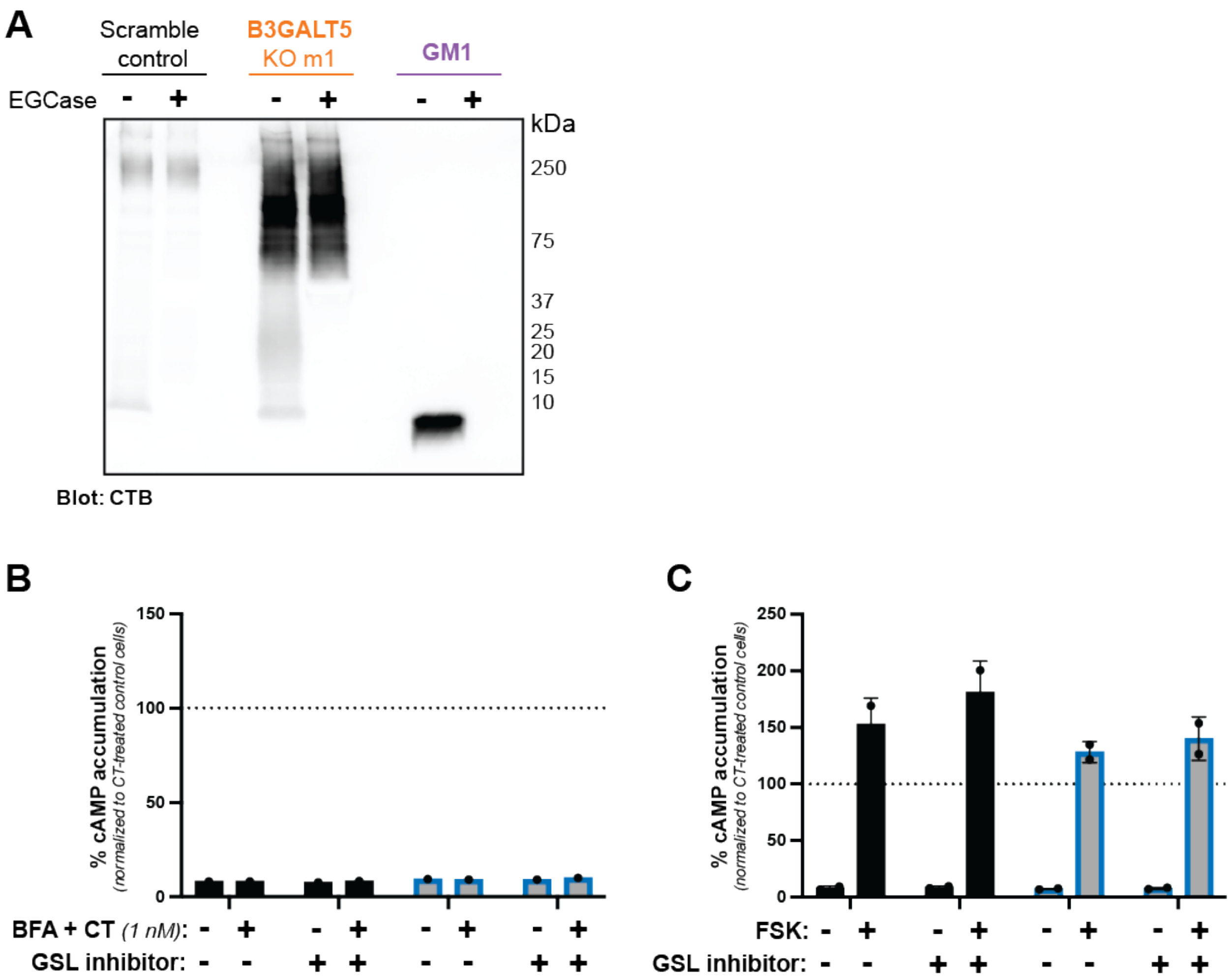


**Supplementary Figure 5. Validation of GSLs as decoy receptors for CT**

(**A**) Lectin blot with CTB-biotin of control and B3GALT5 KO m1 cell lysates, and pure GM1. Samples were treated for 16 h with endoglycoceramidase or a vehicle control. Data shown are a single representative trial of 3 independent biological replicates. (**B**, **C**) Control and B3GNT5 KO+OE cells were treated with P4 inhibitor of glycosphingolipid biosynthesis for 72 h. Cells pretreated with brefeldin A (BFA) or a vehicle control for 0.5 h were incubated for 1.5 h with CT (1 nM) or buffer alone (**B**). Alternately, cells were treated with forskolin (10 μM) for 0.5 h (**C**). Accumulation of cAMP was measured. Data shown are inverse of luminescence values normalized first to the total amount of cells plated for each cell line, then to the signal in CT-treated control cells. Each datapoint is a biological replicate consisting of 3 averaged technical replicates. Panel B represents a single biological replicate. In panel C, error bars indicate mean ± SD of 2 biological replicates. Statistical analysis was by two-way ANOVA with Tukey correction. No significant differences were detected.

## A

| M+Na | Composition | Proposed Glycoform | Scramble Control *Average (±STDEV)* | B3GALT5 KO *Average (±STDEV)* |
|---|---|---|---|---|
| 1141.572 | (Hex)2 (HexNAc)2 (Deoxyhexose)1 | | 6.5% (±0.7%) | 2.6% (±0.3%) |
| 1171.563 | (Hex)3 (HexNAc)2 | | 1.6% (±0.1%) | 0.9% (±0.1%) |
| 1345.672 | (Hex)3 (HexNAc)2 (Deoxyhexose)1 | | 13.7% (±1.6%) | 3.7% (±1.3%) |
| 1375.683 | (Hex)4 (HexNAc)2 | | 2.6% (±0.2%) | 1.1% (±0.8%) |
| 1538.756 | (Hex)6 (HexNAc)1 | | 0.6% (±0.5%) | 1.0% (±0.2%) |
| 1579.783 | (Hex)2 + (Man)3(GlcNAc)2 | | 8.2% (±0.8%) | 7.0% (±0.2%) |
| 1590.798 | (HexNAc)1 (Deoxyhexose)1 + (Man)3(GlcNAc)2 | | 0.5% (±0.0%) | 0.3% (±0.1%) |
| 1742.854 | (Hex)7 (HexNAc)1 | | 0.6% (±0.1%) | 0.5% (±0.4%) |
| 1783.882 | (Hex)3 + (Man)3(GlcNAc)2 | | 17.5% (±1.4%) | 15.4% (±3.8 %) |
| 1835.924 | (HexNAc)2 (Deoxyhexose)1 + (Man)3(GlcNAc)2 | | 1.4% (±0.3%) | 0.8% (±0.1%) |
| 1946.954 | (Hex)8 (HexNAc)1 | | 0.7% (±0.2%) | 1.0% (±0.2%) |
| 1987.982 | (Hex)4 + (Man)3(GlcNAc)2 | | 8.4% (±0.5%) | 10.5% (±0.5%) |
| 2040.022 | (Hex)1 (HexNAc)2 (Deoxyhexose)1 + (Man)3(GlcNAc)2 | | 0.7% (±0.2%) | 0.4% (±0.1%) |
| 2070.029 | (Hex)2 (HexNAc)2 + (Man)3(GlcNAc)2 | | 0.7% (±0.1%) | 1.2% (±0.1%) |
| 2151.049 | (Hex)9 (HexNAc)1 | | 0.6% (±0.1%) | 1.2% (±0.1%) |
| 2156.073 | (Hex)1 (HexNAc)1 (Deoxyhexose)1 (NeuAc)1 + (Man)3(GlcNAc)2 | | 0.9% (±0.3%) | 1.1% (±0.5%) |
| 2192.081 | (Hex)5 + (Man)3(GlcNAc)2 | | 14.8% (±1.6%) | 20.4% (±0.9%) |
| 2203.097 | (Hex)3 (HexNAc)1 (Deoxyhexose)1 + (Man)3(GlcNAc)2 | | 1.1% (±0.0%) | 0.8% (±0.1%) |
| 2244.122 | (Hex)2 (HexNAc)2 (Deoxyhexose)1 + (Man)3(GlcNAc)2 | | 0.8% (±0.1%) | 0.9% (±0.1%) |
| 2274.135 | (Hex)3 (HexNAc)2 + (Man)3(GlcNAc)2 | | 0.6% (±0.1%) | 0.7% (±0.3%) |
| 2285.149 | (Hex)1 (HexNAc)3 (Deoxyhexose)1 + (Man)3(GlcNAc)2 | | 0.7% (±0.1%) | 0.5% (±0.1%) |
| 2390.182 | (Hex)3 (HexNAc)1 (NeuAc)1 + (Man)3(GlcNAc)2 | | 0.3% (±0.0%) | 0.6% (±0.0%) |
| 2396.181 | (Hex)6 + (Man)3(GlcNAc)2 | | 7.6% (±0.8%) | 12.6% (±0.6%) |
| 2401.194 | (Hex)1 (HexNAc)2 (Deoxyhexose)1 (NeuAc)1 + (Man)3(GlcNAc)2 | | 0.4% (±0.0%) | 0.8% (±0.0%) |
| 2418.209 | (Hex)2 (HexNAc)2 (Deoxyhexose)2 + (Man)3(GlcNAc)2 | | 0.5% (±0.1%) | 0.6% (±0.1%) |
| 2448.224 | (Hex)3 (HexNAc)2 (Deoxyhexose)1 + (Man)3(GlcNAc)2 | | 0.3% (±0.2%) | 0.8% (±0.1%) |
| 2459.238 | (Hex)1 (HexNAc)3 (Deoxyhexose)2 + (Man)3(GlcNAc)2 | | 0.5% (±0.1%) | 0.6% (±0.1%) |
| 2489.25 | (Hex)2 (HexNAc)3 (Deoxyhexose)1 + (Man)3(GlcNAc)2 | | 0.9% (±0.1%) | 0.7% (±0.1%) |
| 2592.304 | (Hex)2 (HexNAc)2 (Deoxyhexose)3 + (Man)3(GlcNAc)2 | | 0.5% (±0.1%) | 0.6% (±0.0%) |
| 2600.282 | (Hex)7 + (Man)3(GlcNAc)2 | | 1.1% (±0.1%) | 1.7% (±0.0%) |
| 2652.326 | (Hex)4 (HexNAc)2 (Deoxyhexose)1 + (Man)3(GlcNAc)2 | | 0.3% (±0.1%) | 0.6% (±0.1%) |
| 2663.337 | (Hex)2 (HexNAc)3 (Deoxyhexose)2 + (Man)3(GlcNAc)2 | | 1.1% (±0.2%) | 1.2% (±0.2%) |
| 2779.385 | (Hex)2 (HexNAc)2 (Deoxyhexose)2 (NeuAc)1 + (Man)3(GlcNAc)2 | | 0.2% (±0.1%) | 0.4% (±0.0%) |
| 2837.425 | (Hex)2 (HexNAc)3 (Deoxyhexose)3 + (Man)3(GlcNAc)2 | | 1.5% (±0.3%) | 2.2% (±0.4%) |
| 2850.421 | (Hex)2 (HexNAc)3 (Deoxyhexose)1 (NeuAc)1 + (Man)3(GlcNAc)2 | | 0.2% (±0.1%) | 0.6% (±0.1%) |
| 3024.511 | (Hex)2 (HexNAc)3 (Deoxyhexose)2 (NeuAc)1 + (Man)3(GlcNAc)2 | | 0.5% (±0.1%) | 1.5% (±0.1%) |
| 3198.601 | (Hex)2 (HexNAc)3 (Deoxyhexose)3 (NeuAc)1 + (Man)3(GlcNAc)2 | | 0.1% (±0.0%) | 0.5% (±0.2%) |
| 3215.613 | (Hex)3 (HexNAc)3 (Deoxyhexose)4 + (Man)3(GlcNAc)2 | | 0.2% (±0.0%) | 0.3% (±0.0%) |
| 3228.601 | (Hex)3 (HexNAc)3 (Deoxyhexose)2 (NeuAc)1 + (Man)3(GlcNAc)2 | | 0.3% (±0.1%) | 1.2% (±0.7%) |
| 3286.648 | (Hex)3 (HexNAc)4 (Deoxyhexose)3 + (Man)3(GlcNAc)2 | | 0.3% (±0.1%) | 0.3% (±0.1%) |

**Note:** Glycoforms shown represent only one of multiple possible isomers

## B

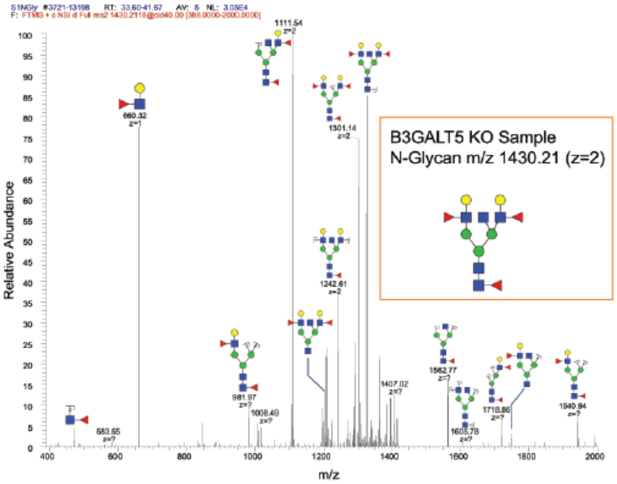


## C

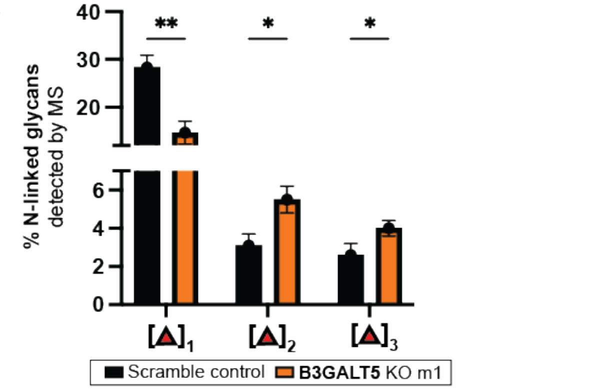


**Supplementary Figure 6. B3GALT5 KO cells exhibit increased fucosylation on N-linked glycoproteins**
(**A**) N-linked glycoforms detected in control and B3GALT5 KO cells by LC-MS/MS analysis. Quantification is based on triplicate analysis. (**B**) Example MS/MS of a complex, fucosylated N-linked glycan detected in B3GALT5 KO cells confirming placement of fucose. (**C**) Bar graphs showing the relative enrichment of mono- vs di- vs tri-fucosylated N-linked glycans detected by LC-MS/MS analysis of control and B3GALT5 KO cells. Statistical analysis was performed by t-test with Holm-Šídák correction.

**A**

| M+Na | Composition | Proposed Glycoform | Scramble Control Average (±STDEV) | B3GALT5 KO Average (±STDEV) |
|---|---|---|---|---|
| 534.2875 | $(Hex)_1$ $(HexNAc)_1$ | | 64.7% (±5.0%) | 4.3% (±1.4%) |
| 650.3345 | $(Hex)_1$ $(NeuAc)_1$ | | 3.6% (±2.0%) | 3.7% (±1.0%) |
| 779.4017 | $(Hex)_1$ $(HexNAc)_2$ | | 9.7% (±2.2%) | 1.6% (±1.0%) |
| 895.4597 | $(Hex)_1$ $(HexNAc)_1$ $(NeuAc)_1$ | | 8.4% (±3.2%) | 16.1% (±1.3%) |
| 983.5113 | $(Hex)_2$ $(HexNAc)_2$ | | 12.6% (±2.8%) | 3.4% (±1.3%) |
| 1157.601 | $(Hex)_2$ $(HexNAc)_2$ $(Deoxyhexose)_1$ | | 0.3% (±0.2%) | 1.8% (±2.1%) |
| 1256.634 | $(Hex)_1$ $(HexNAc)_1$ $(NeuAc)_2$ | | 0.6% (±0.6%) | 19.7% (±1.5%) |
| 1344.685 | $(Hex)_2$ $(HexNAc)_2$ $(NeuAc)_1$ | | N.D. | 7.0% (±0.3%) |
| 1518.776 | $(Hex)_2$ $(HexNAc)_2$ $(Deoxyhexose)_1$ $(NeuAc)_1$ | | N.D. | 10.6% (±0.6%) |
| 1606.829 | $(Hex)_3$ $(HexNAc)_3$ $(Deoxyhexose)_1$ | | N.D. | 1.3% (±0.1%) |
| 1705.861 | $(Hex)_2$ $(HexNAc)_2$ $(NeuAc)_2$ | | N.D. | 2.2% (±0.1%) |
| 1879.949 | $(Hex)_2$ $(HexNAc)_2$ $(Deoxyhexose)_1$ $(NeuAc)_2$ | | N.D. | 14.2% (±1.6%) |
| 1881.956 | $(Hex)_4$ $(HexNAc)_4$ | | N.D. | 7.2% (±0.9%) |
| 1968.001 | $(Hex)_3$ $(HexNAc)_3$ $(Deoxyhexose)_1$ $(NeuAc)_1$ | | N.D. | 1.2% (±0.1%) |
| 2142.091 | $(Hex)_3$ $(HexNAc)_3$ $(Deoxyhexose)_2$ $(NeuAc)_1$ | | N.D. | 1.9% (±0.1%) |
| 2417.229 | $(Hex)_4$ $(HexNAc)_4$ $(Deoxyhexose)_1$ $(NeuAc)_1$ | | N.D. | 1.3% (±0.1%) |
| 2503.265 | $(Hex)_3$ $(HexNAc)_3$ $(Deoxyhexose)_2$ $(NeuAc)_2$ | | N.D. | 2.4% (±0.3%) |

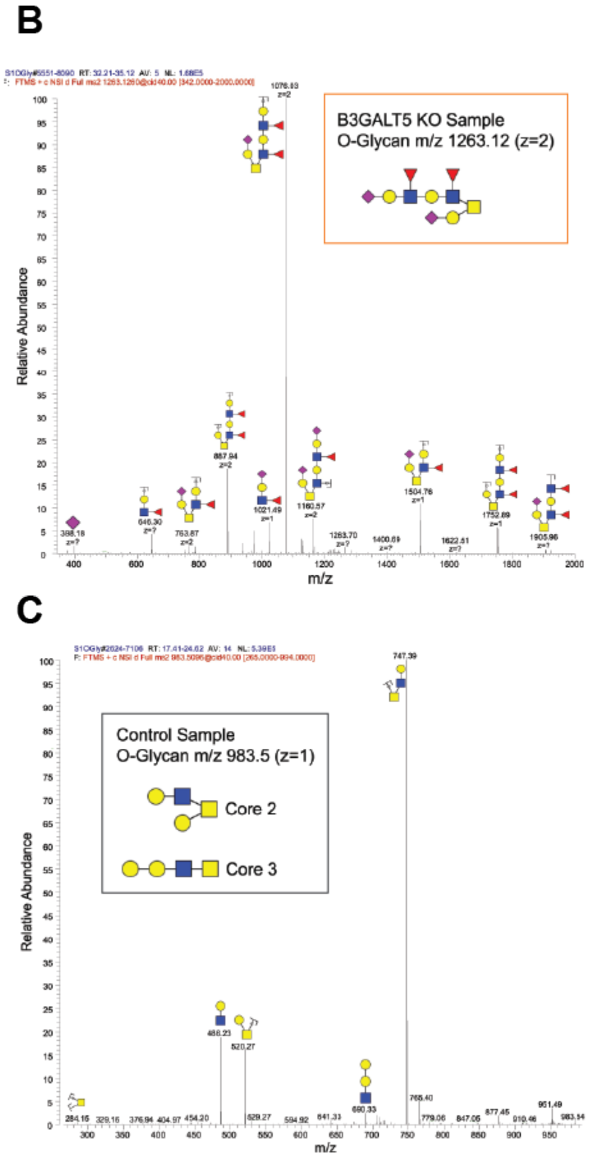


**Supplementary Figure 7. B3GALT5 KO cells exhibit increased fucosylation on O-linked glycoproteins**
(**A**) O-linked glycoforms detected in control and B3GALT5 KO cells by LC-MS/MS analysis. Quantification is based on triplicate analysis. (**B**) Example MS/MS of a fucosylated O-linked glycan detected in B3GALT5 KO cells. (**C**) Example MS/MS of core 2 O-linked glycan detected in B3GALT5 KO cells confirming structure.

**A**

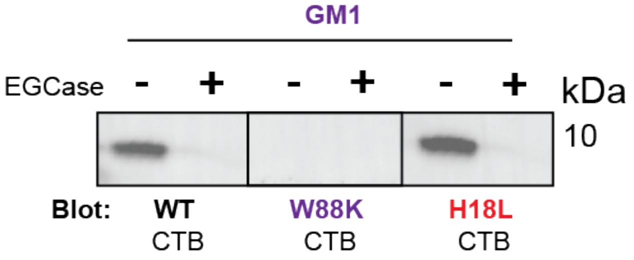


**Supplementary Figure 8. CTB binding to GM1 depends on the canonical glycan binding pocket**
GM1 was treated with EGCase or vehicle, then detected by lectin blot using WT CTB-biotin, W88K CTB-biotin, and H18L CTB-biotin. Data presented are a single replicate.

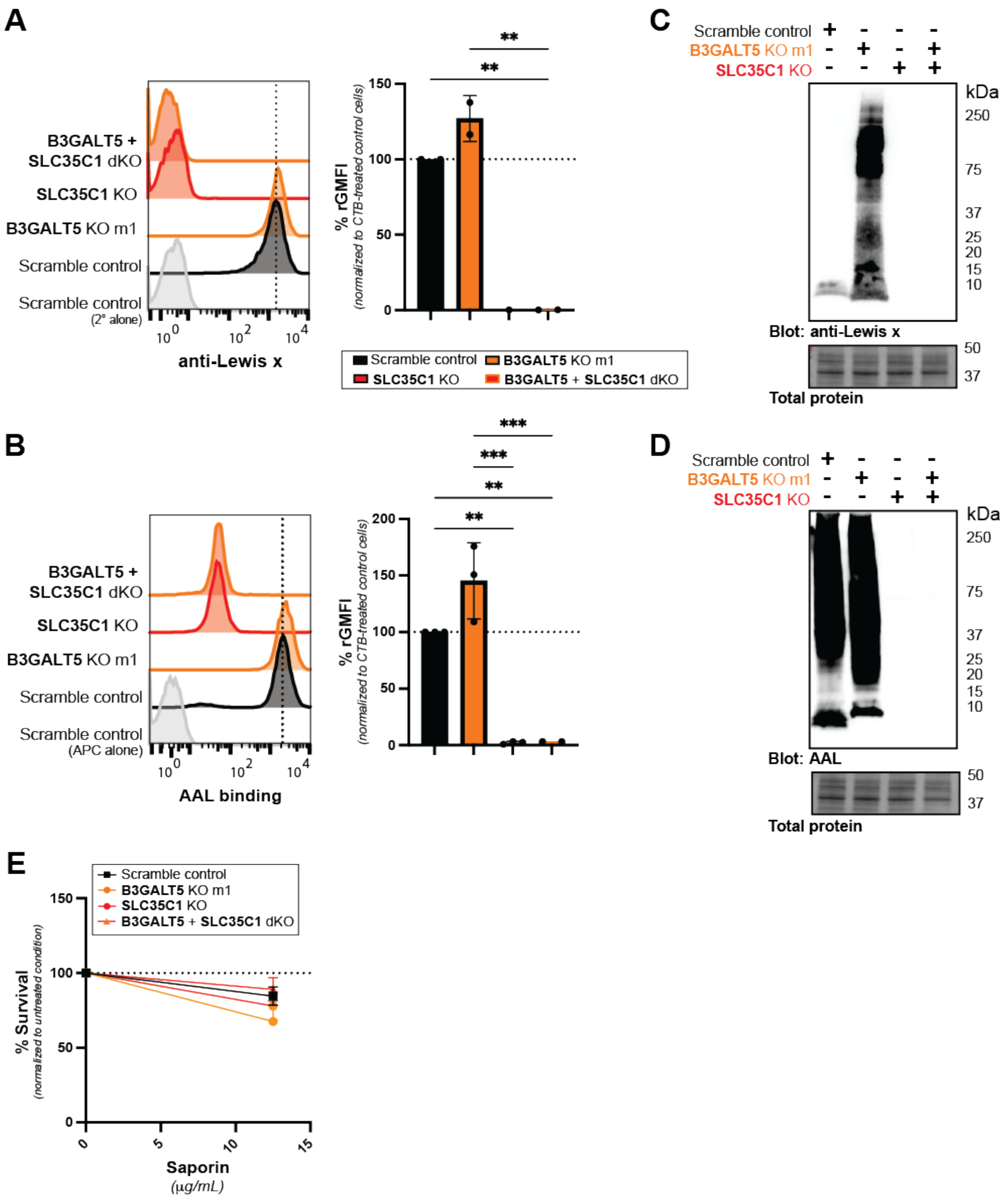


**Supplementary Figure 9. Validation of SLC35C1 KO cell lines**

Representative histograms from flow cytometry analyses of anti-Le$^{x}$ antibody (**A**) and AAL (**B**) binding to surfaces of control, B3GALT5 KO m1, SLC35C1 KO, and B3GALT5+SLC35C1 dKO cells. Quantification of gMFIs from 1 – 3 biological replicates are normalized to the maximum signal in control cells. Lysates from control, B3GALT5 KO, SLC35C1 KO and B3GALT5+SLC35C1 double KO cells were analyzed by immunoblot probing with anti-Le$^{x}$ antibody (**C**) or lectin blot probing with AAL (**D**). Data shown are a single biological replicate. (**E**) Control, B3GALT5 KO m1, SLC35C1 KO and B3GALT5+SLC35C1 dKO cells were incubated for 72 h with unconjugated saporin (12.5 μg/mL). Survival data shown are luminescence values normalized to the signal from the untreated condition for each cell type. Each datapoint is a biological replicate consisting of 3 averaged technical replicates. Error bars indicate mean ± SD of 2 biological replicates. (**F**) Cells pretreated with brefeldin A (BFA) or a vehicle control for 0.5 h were incubated for 1.5 h with CT (1 nM) or buffer alone. Accumulation of cAMP was measured. Data shown are inverse of luminescence values normalized first to the total amount of cells plated for each cell line, then to the signal in CT-treated control cells. Data presented are a single biological replicate consisting of 3 averaged technical replicates. (**G**) Cells were treated with forskolin (10 μM) for 0.5 h and then analyzed as in panel F. Data presented are represent two biological replicates.

### Supplementary Table 1. Sequence Validation of KO cell lines

| | | Nucleic Acid Sequence | InDel |
|---|---|---|---|
| **B3GALT5 KO sgRNA** | | CATCGCTTTTGTCCTCAGG | |
| **B3GALT5 WT gene** | | GACCATGATGGGCATAGAATGGGTCCATCGCTTTTGTCCTCAGGCGGCGTTTGTGATGAAAACAGACT | 0 |
| B3GALT5 KO m1 | Allele 1 | GACCATGATGGGCATAGAATGGGTCCATCGCTTTTGTCCTCTAGGCGGCGTTTGTGATGAAAACAGACT | 1 |
|  | Allele 2 | GACCATGATGGGCATA--------------------------------------------------------------------------------------------------------------AAC | -110 |
| B3GALT5 KO m2 | Allele 1 | GACCATGATGGGCATAGAATGGGTCCATCGCTTTTGT-------CGGCGTTTGTGATGAAAACAGACT | -7 |
| | Allele 2 | GAAGACCATGATGGGCATAGAATGGGTCCATCGCTTTTGTCCTC 223 BP insertion AGGCGGCGTTTGTGATGAAAACAGACT | 223 |
| **B3GNT5 KO sgRNA** | | GGATTGGTCGTGTTCATCG | |
| **B3GNT5 WT gene** | | AACAAATTGGTGTTCAAGACTTTTGGATTGGTCGTGTTCATCGTGGTGCCCCTCCCATTAGAGATAAA | 0 |
| B3GNT5 KO m1 | Allele 1 | AACAAATTGGTGTTCAAGACTTTTGGATTGGTCGTGTTCAATCGTGGTGCCCCTCCCATTAGAGATAAA | 1 |
| | Allele 2 | AACAAATTGGTGTTCAAGACTTTTGGATTGGTCGTGTT--ATCGTGGTGCCCCTCCCATTAGAGATAAA | -2 |
| B3GNT5 KO m2 | Allele 1 | AACAAATTGGTGTTCAAGACTTTTGGATTGGTCGTGTTCA-CGTGGTGCCCCTCCCATTAGAGATAAA | -1 |
| | Allele 2 | AACAAATTGGTGTTCAAGACTTTTGGATTGGTCGTGTTCATTCGTGGTGCCCCTCCCATTAGAGATAAA | 1 |
| B3GNT5 KO in | Allele 1 | AACAAATTGGTGTTCAAGACTTTTGGATTGGTCGTGTTCAATCGTGGTGCCCCTCCCATTAGAGATAAA | 1 |
| B3GALT5 KO m1 | Allele 2 | AACAAATTGGTGTTCAAGACTTTTGGATTGGTCGTGTTCA------------TCCCATTAGAGATAAA | -12 |
| **SLC35C1 KO sgRNA** | | ACCACGAAGGTGCTCCCGG | |
| **SLC35C1 WT gene** | | CTGTGTCTCGCTCAACGCCATCTACACCACGAAGGTGCTCCCGGCGGTGGACGGCAGCATCTGGCGCC | 0 |
| SLC35C1 KO | Allele 1 | CTGTGTCTCGCTCAACGCCATCTACACCACGAAGGTGCT--CGGCGGTGGACGGCAGCATCTGGCGCC | -2 |
|  | Allele 2 | CTGTGTCTCGCTCAACGCCATCTACACCACGA---------CGGCGGTGGACGGCAGCATCTGGCGCC | -9 |
| SLC35C1 KO in B3GALT5 KO m1 | Allele 1 | CTGTGTCTCGCTCAACGCCATCTACACCACGA------------CGGTGGACGGCAGCATCTGGCGCC | -12 |
